# Learning by forgetting: A computational model of insect brain

**DOI:** 10.64898/2026.04.21.719789

**Authors:** Koichiro Yamauchi, Aditya Gyanprakash Nirmale

**Affiliations:** Department of Computer Science, Chubu University, Kasugai, Matsumoto-cho 1200, Japan; Department of Electrical Engineering, Indian Institute of Technology Bombay, Powai, Mumbai – 400076, India

## Abstract

In this study, resource-constrained learning methods were developed as a model for the learning behavior of the fly brain, specifically the mushroom body. Recent research on the mushroom bodies of flies shows that unfamiliar odors activate certain output neurons (MBONs); however, these effects are rapidly suppressed upon repeated exposure to the same odor. Such MBON behaviors appear to reflect odor learning. We investigated how flies continue learning about odors throughout their lives despite their small brains. Researchers have suggested that learning about new odors can help flies forget existing memories. Therefore, we hypothesized that the main reason for continual learning is that it serves as a strategy for forgetting. To test the validity of this hypothesis, we designed three models using a kernel perceptron. This approach is suitable for estimating ongoing learning capacity within a budget. According to the results of computer simulations and theoretical analysis, the model demonstrated the importance of forgetting mechanisms for two reasons: first, to prepare for subsequent learning sessions, and second, to reduce the negative effects of deleting memories.

**Author summary:** Drosophila mushroom body output neurons (MBONs) in the *α*’3 compartment of the fruit fly brain are highly activated by novel odors, and their activation triggers alerting behavior. Interestingly, these specific neurons react only to unfamiliar odor information, suggesting they constantly undergo incremental learning of new odors. This study was aimed at constructing three incremental learning models of the MBON *α*’3 neurons. Although there have been numerous studies on complex circuit designs to reproduce activation waveforms, herein we constructed a fundamental learning model based on a kernelized learning method. Since kernelized learning models interpret Hebbian learning as the addition or subtraction of kernel functions, the model is easy to analyze theoretically. Consequently, we conclude that the forgetting property observed in the MBON *α*’3 neurons is essential for reducing error when learning occurs within a brain of limited capacity.

## Introduction

In insects, mushroom bodies are responsible for memory and learning [1]. The nerve center of *Drosophila* is a mushroom body that consists mainly of Kenyon cells (KCs) [1], mushroom body output neurons (MBONs), and dopaminergic neurons (DANs). Some DANs are well known to encode the predicted reward of the current situation, and their output signals affect the connections between KCs and MBONs. Moreover, DANs control the learning of MBONs, which in turn send output signals to other areas of the insect brain.

Odors are sensed by olfactory sensory neurons in the antennae of *Drosophila*, and the gathered information is distributed to KCs by projection neurons (PNs). The connections between PNs and KCs appear to be random and do not exhibit plasticity. Approximately 2000 KCs exist in this region, and each KC neuron represents a sensory input. KCs send their axonal fibers to MBONs, which are divided into 15 compartments, each with its own specific function. Plasticity is observed in the connections between KCs and MBONs. The connections between KCs and MBONs are adjusted according to the reward signals from DANs. The main role of DANs is to represent predicted reward signals. Additionally, DANs represent the novelty of current stimuli.

Hattori et al. [2] investigated the MBONs in the *α*’3 compartments of *Drosophila* by activating them with novel odors. When the MBON *α*’3 neurons were activated, *Drosophila* exhibited an alerting behavior. This novel order-evoked MBON activity was quickly suppressed by repeated stimulation with the same odor, suggesting that *Drosophila* rapidly learns to recognize a novel odor upon repeated stimulation, which renders the odor progressively more familiar. Thus, *Drosophila* can engage in incremental learning of novel odors, and odor perception is critical to their life. The estimated novelty-detection capacity translates into 10–20 orthogonal feature vectors and implies that MBONs can store 10–20 memory units. However, 10–20 memory units may be significantly less for *Drosophila* to survive for a month. This observation raises a fundamental question: how does the fruit fly maintain continuous learning despite such limited neural resources?

To develop technological breakthroughs, imitation of biological systems is sometimes valuable. This study is focused on the *Drosophila* brain, which contains relatively few neurons. Due to limited resources, performing incremental learning is challenging. Therefore, we try to gain insights into how small brains learn by studying the *Drosophila* brain. This study develops a learning model for MBONs and DANs based on a kernel perceptron. The kernel perceptron framework is particularly suitable for mistake-bound analyses, which allow for a formal evaluation of learning efficiency. Using this approach, we try to characterize the learning properties of *Drosophila* within a mistake-bound paradigm. For analysis, Forgetron [3] is considered the base model because it reduces the weight parameters during incremental learning, such that the model fits the learning strategy of MBONs. However, the mistake-bound model assumes that the learning machine forgets the oldest memory in each round. The purpose of forgetting is to decrease dependence on each memory and reduce the adverse effects of removing the oldest memory. Conversely, forgetting one memory is a way to make room for new learning.

Furthermore, a simulation study is performed using Forgetron and a modified version of Forgetron in changing environments. The observed results assist in determining the effects of forgetting. This study builds on and extends the preliminary results presented in our previous research [4]. To clarify the novel contributions of this study, the main improvements over previous studies are as follows:

1. **Clarification of Simulation Results:** Simulation settings are refined, and the results are analyzed in greater detail to better demonstrate the effects of the forgetting mechanism.
2. **Theoretical Analysis of the Least Recently Used** (**LRU) Model:** For the LRU strategy (Model 3), a detailed theoretical analysis is performed using Markov chains and the ergodic theorem, thereby establishing the performance advantage of the LRU model.
3. **Sensitivity Analysis of the Shrinkage Coefficient (***ϕ*): Sensitivity analysis of the shrinkage coefficient *ϕ* is conducted to verify the effect of the mistake-bound model. By detailing the minimization of *cumulative error*, we theoretically demonstrate that an optimal forgetting rate exists for MBONs to minimize the number of prediction errors.

## Results

### Kernelized incremental learning on a budget as a model of MBON *α*^′^ 3 neurons

In this study, we develop a model of the MBONs in the *α*’3 compartment of *Drosophila*. The *α*’3 compartment contains a single MBON layer along with associated dopamine neurons (PPL1*α*’-3), which regulate alerting behaviors in response to novel odors. The memory capacity of this compartment depends on the number of orthogonal vectors in KCs, typically ranging from 10 to 20. Hattori et al. [2] reported that learning a novel odor reduces existing memory in MBON *α*’3 neurons. This finding suggests that acquiring new odor memories leads to the removal of part of the existing memory, which in turn facilitates the learning of additional new odors in the future.

A key constraint of MBONs is the limited number of neurons available for memory storage. As new stimuli are encountered sequentially, the system must learn and use these memories within a fixed resource budget. If the memory capacity becomes saturated and no further forgetting occurs, the system may fail to adapt to new situations. To solve this problem, forgetting a portion of existing memory is necessary; however, this introduces an additional risk. Specifically, forgetting can (i) cause sudden changes in knowledge structure owing to abrupt memory pruning, which may interfere with existing knowledge, and (ii) result in the loss of information that could become important again in the future, thereby reducing adaptability.

To systematically investigate these trade-offs, we propose three models, each corresponding to one of the following challenges:

Model 1 addresses the need to forget parts of memory to create space for new learning (adaptability to new situations).

Model 2 focuses on minimizing the negative impact of the interference caused by abrupt forgetting (stability of existing knowledge).

Model 3 considers the risk of being unable to recall previously forgotten knowledge when it becomes relevant (long-term adaptability).

By comparing these models, we aim to clarify how different forgetting mechanisms affect the balance between adaptability and memory retention under resource constraints. Similar to existing models [5, 6], the proposed MBON model uses Hebbian learning guided by DANs. The main difference is that the proposed model is based on the kernel method and is designed to evaluate incremental learning under a fixed memory budget. Unlike previous models, the designed approach does not use reward signals because learning is assumed to be driven by the free-energy principle. Therefore, the model does not require a reinforcement learning environment.

### Preliminary: Kernelized representation of MBONs

Although the *α*’3 compartment is highly likely to employ anti-Hebbian learning, its functionality can also be represented by a Hebbian learning model. Therefore, a novelty detector can be constructed using an active neuron that is inhibited by a Hebbian learning-based odor detector. This study uses kernelized Hebbian learning to represent the function in a simplified manner (more details are provided in the **Validity of analyzing Anti-Hebbian learning as Hebbian learning** section). As discussed in **the Introduction**, approximately 2000 KCs represent random feature vectors of the sensory inputs received by the MBONs through synaptic connections. An odor activates approximately 5–10% of the 2000 KCs. These sparse feature vectors are nearly orthogonal to one another. Thus, sensory inputs are projected onto a higher-dimensional space through an orthogonal transformation, and MBONs process high-dimensional feature vectors for classification or regression. A high-dimensional feature mapping 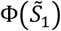 and 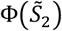, obtained from input vectors 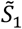 and 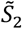 using a specialized nonlinear function Φ(·), has a dot product 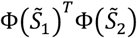 that is represented by a reproducing the kernel function 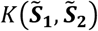 [7]. One such kernel function is the Gaussian kernel function defined as follows:

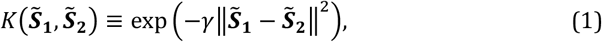

where *γ* is a coefficient that depends on the nonlinear function Φ(·). Using this analogy, we modify the multiplication involving the KC output vector 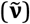. The synaptic weight matrix (***W***_*mbon*_) is approximated using the kernel function output as follows:

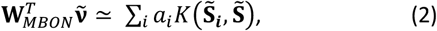

where 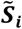 the *i*-th support input vector memorized by MBONs, and a_*i*_ is the output vector corresponding to 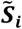. Hebbian and anti-Hebbian learning can be represented as adding or subtracting a target vector 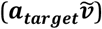 to or from *W*_*MBON*_. In the kernelized algorithm, this corresponds to adding or pruning the kernel function 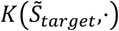. We approximate this process using a kernel perceptron with a limited number of kernels bounded by capacity (budget) *B* (Figure 1).

**Fig. 1.**
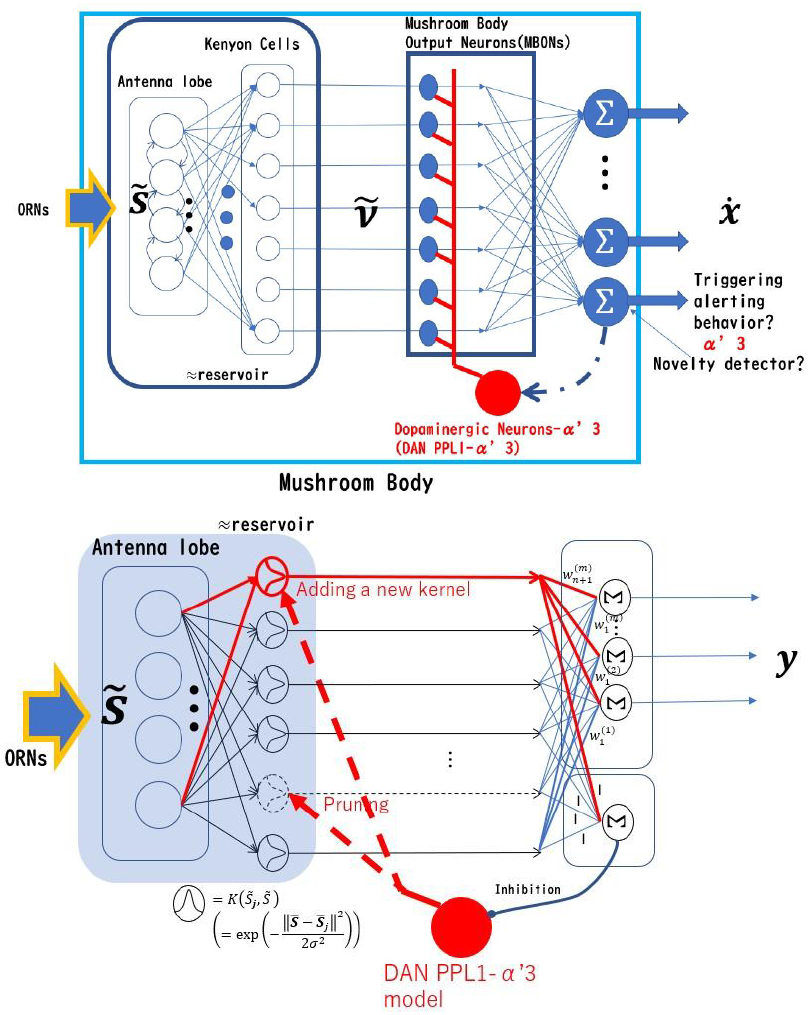
MBON and its kernelized model. Upper: Schematic of the mushroom body output neuron *α*’3. Lower: Kernelized MBON model.

### Shared structure of the three models: Supervised kernel learning model for the mushroom body *α*^′^ 3

This study develops three MBON *α*’3 learning models that differ primarily in their pruning strategies. In this section, we describe the shared structure of the three models. Assume that the model learns 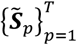 in an online manner, where 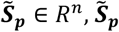 represents a generalized current sensory input vector. The MBON predicts an output vector y ∈ R^*m*^. Notably, under ordinary circumstances, y should be 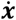; however, in this study, we set y as the class label to focus on MBON learning ability under a fixed budget. 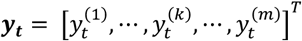, where 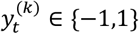.

The model output at time *t* is given by

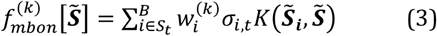

where *B* is the upper bound on the number of hidden units, and *S*_*t*_ is the support set ^1^ at time *t. σ*_*i,t*_ is the decay (forgetting) ratio, where *σ*_*i,t*_ ≤ 1. At each round, *σ*_*i,t*_is updated according to *σ*_*i,t*+1_ = *ϕ*_*t*_*σ*_*i,t*_, where *ϕ*_*t*_ ≤ 1. If the decay ratio *ϕ*_*t*_ is significantly large, it can degrade ***f*** _*mbon*_ and reduce its accuracy. However, pruning old kernels without forgetting them (shrinking) also affects ***f*** _*mbon*_.

Learning samples are presented individually, and the model alternates between inference and learning, as shown in Figure 2. In each round, if at least one of outputs 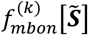 for *k* = 1, …, *m* is incorrect, a new unit (kernel) is added. If the total number of hidden units (kernel functions) exceeds *B*, one unit is removed. For example, the oldest unit is removed (Algorithm 1). Notably, the oldest unit *j* has the smallest *σ*_*i,t*_ because, for all *i* ∈ *S*_*t*_, *σ*_*i,t*_ shrinks according to *σ*_*i,t*+1_ = *ϕ*_*t*_*σ*_*i,t*_ in each round. Thus, the hidden unit with the smallest *σ*_*j,t*_ is removed. Accordingly, the MBON model maintains the total number of hidden units at *B*. Performance is evaluated using cumulative error, as shown in Fig. 2. Note that the weight shrinkage *σ*_*i,t*+1_ = *ϕ*_*t*_*σ*_*i,t*_ is executed only when at least one of the outputs 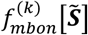 for *k* = 1, …, *m* is incorrect. Thus, weight shrinking is omitted when the outputs are correct, consistent with the MBON*α*^′^3 behaviors reported by [2]. In this framework, the dopaminergic neurons PPL1-*α*^′^3 — primarily activated by novel odors — regulate the weight adaptation of the MBON.

**Fig. 2.**
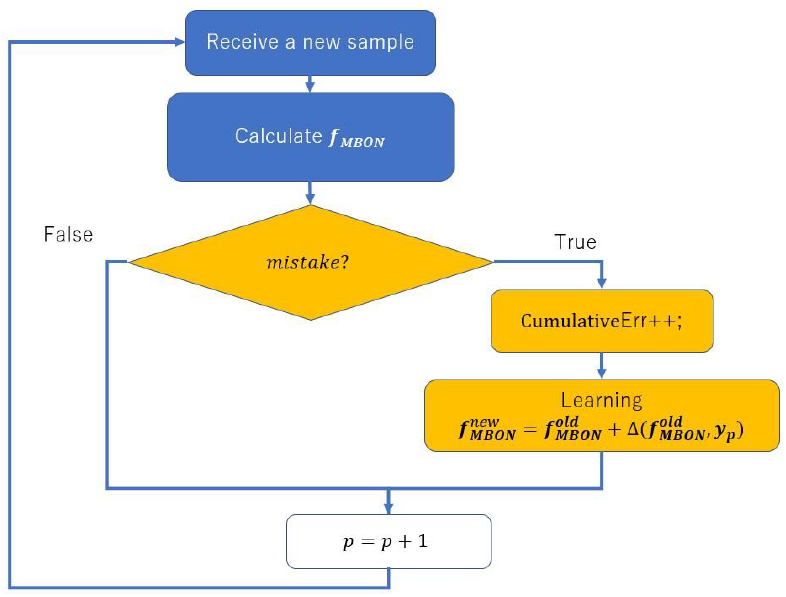
Inference and learning procedure.

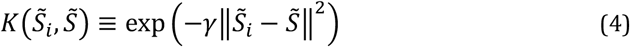

#### Algorithm 1

Kernelized MBON learning Model 1

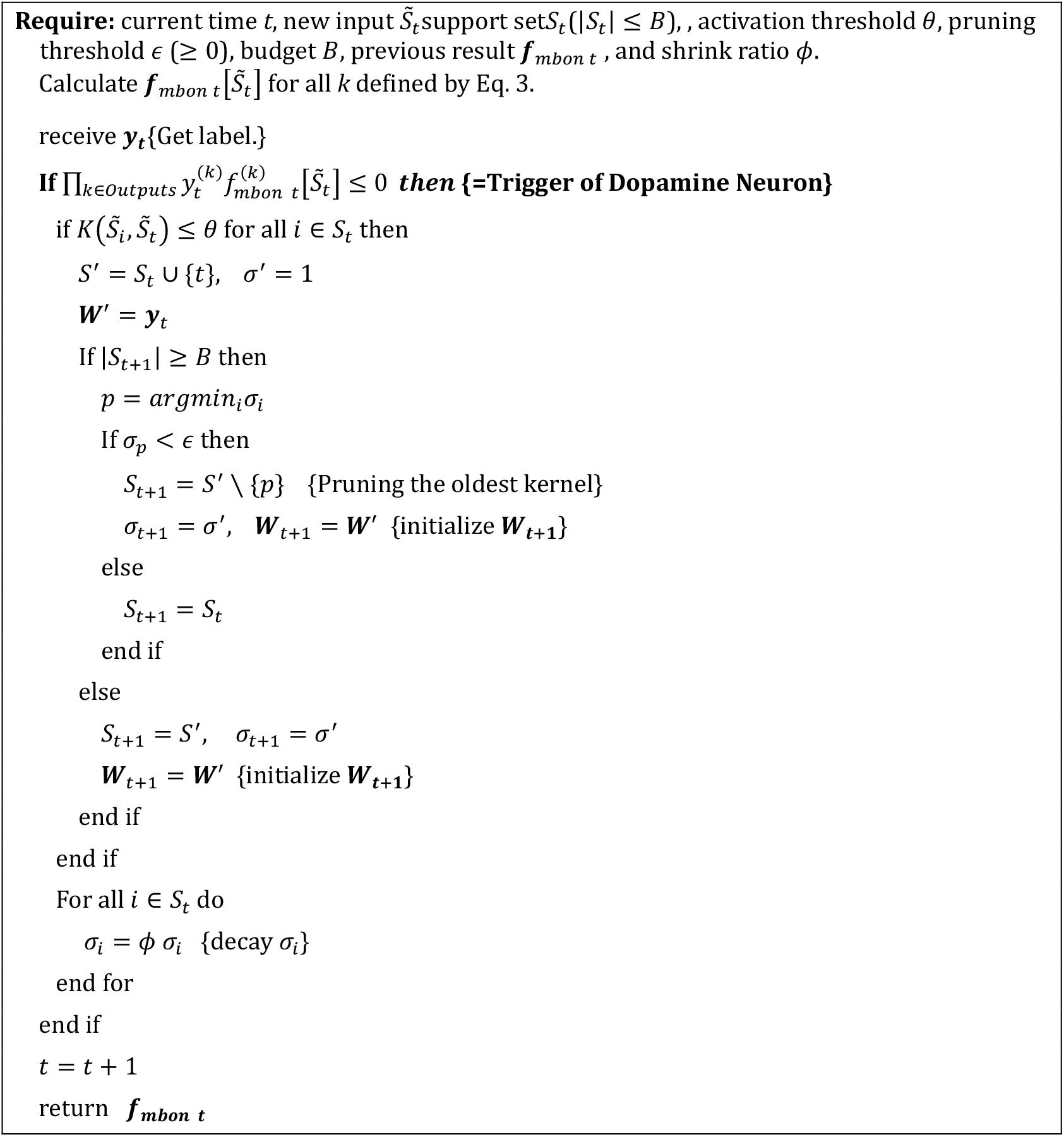

#### Algorithm 2

Kernelized MBON learning Model 2: Modified Forgetron [3]

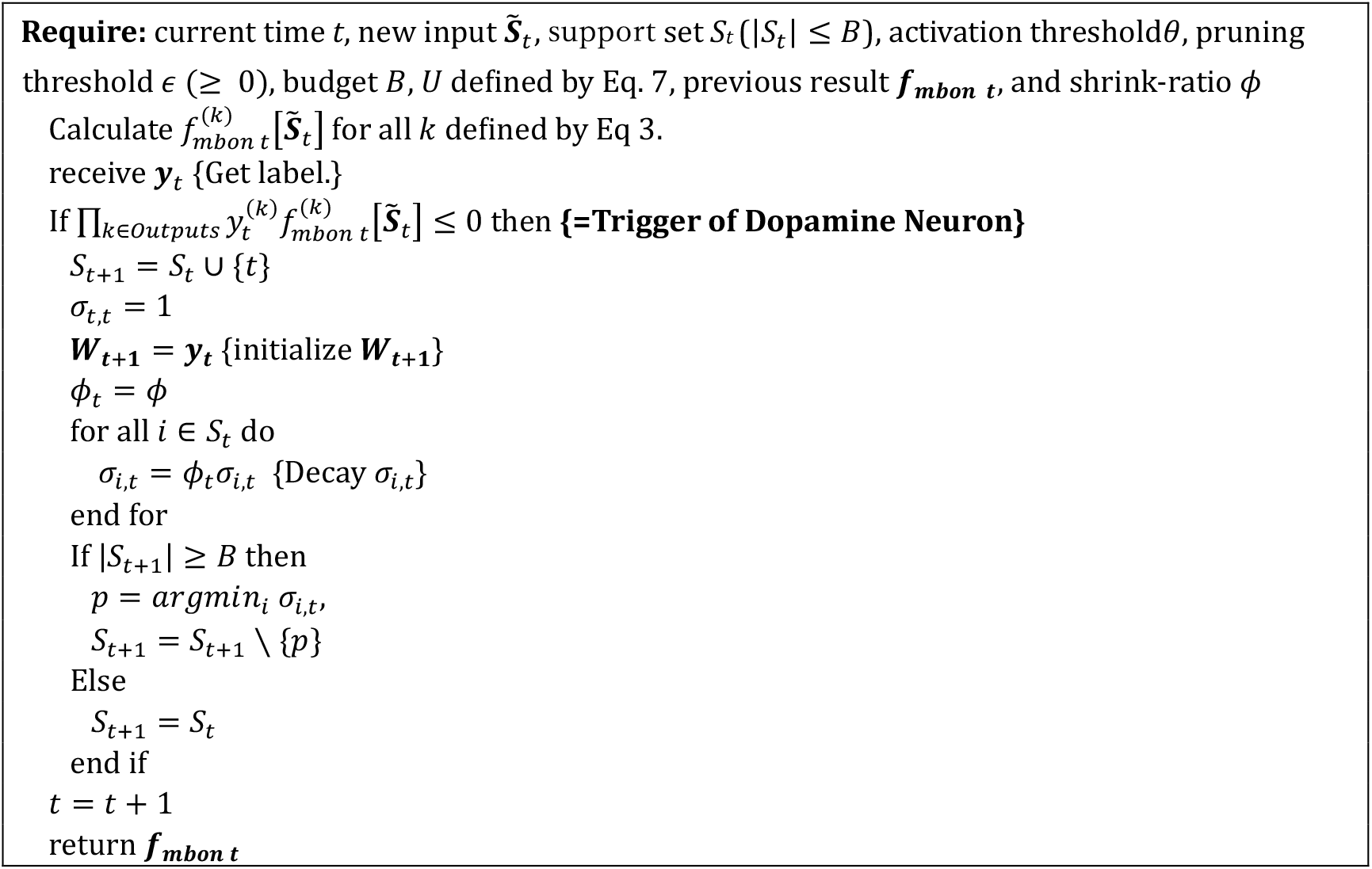

#### Algorithm 3

Kernelized MBON learning Model 3: Least recently used (LRU)

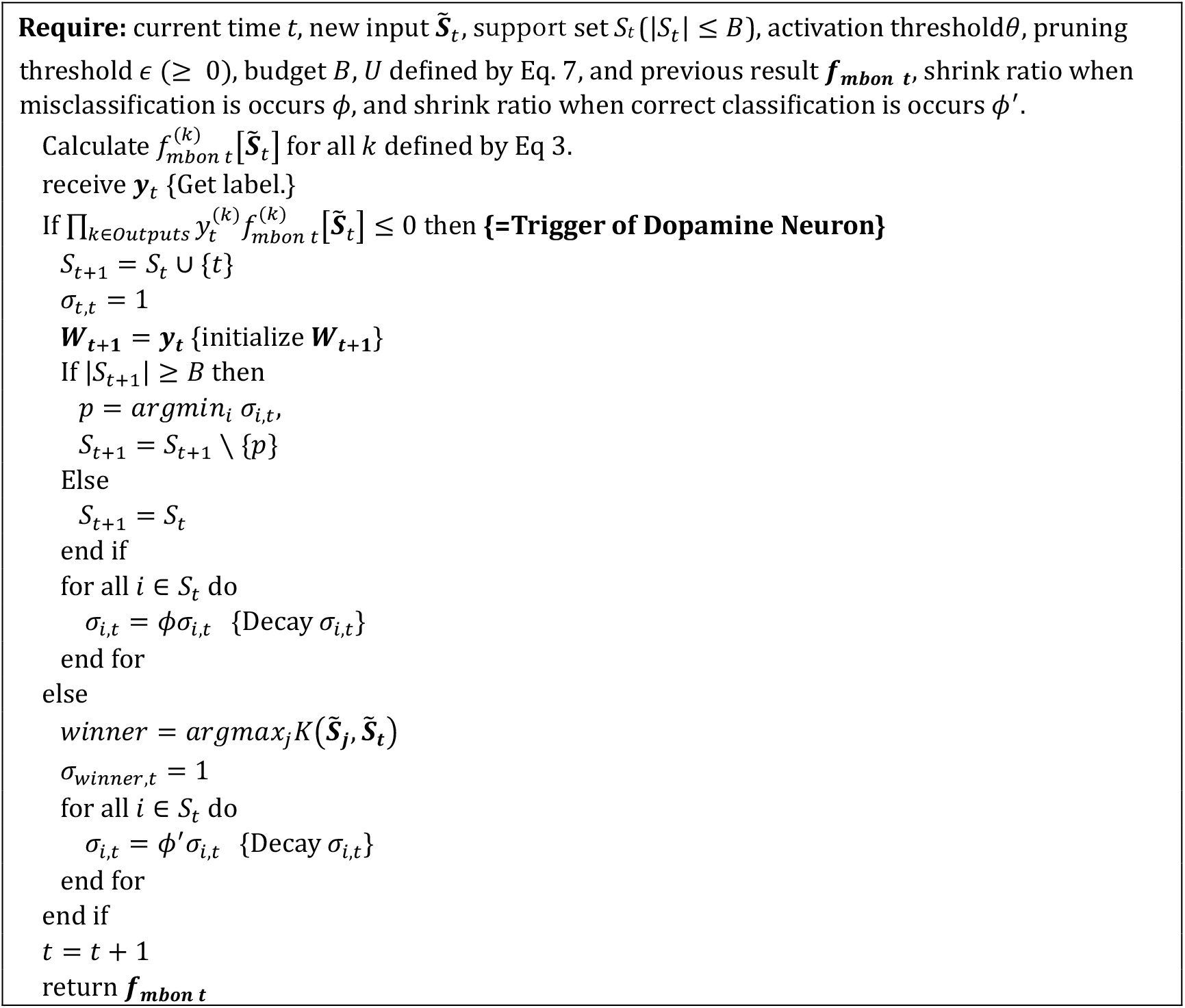

### Three MBON models

#### Model 1

The first model removes the oldest kernel whose *σ*_*i*_is less than *θ* when |*S*_*t*_| ≥ *B*. Thus, if *σ*_*i*_ of the oldest kernel is larger than *θ*, the MBON model does not learn to recognize the current input. Therefore, if *Drosophila* frequently encounters a novel sample, the MBON model often ignores it. The pseudocode is presented in Algorithm 1.

This model detects the oldest kernel via weight shrinking and removes it to create space for learning new samples. However, removing the oldest kernel may cause a significant increase in cumulative error if the kernel remains important for ongoing tasks. Therefore, the threshold *θ* should be set appropriately to balance the trade-off between adaptability to new situations and the stability of existing knowledge.

#### Model 2: Forgetron

We designed a model based on a slightly modified Forgetron [3]. The Forgetron is a kernel perceptron learning machine [8-12] that introduces a weight-shrinking (forgetting) mechanism. The Forgetron removes the oldest kernel whose *σ*_*i*_ is the smallest in *S*_*t*_. Although the Forgetron does not ignore any novel samples, removing the oldest kernel increases cumulative error. The basic Forgetron [3] selects Φ_*t*_ to balance these effects.

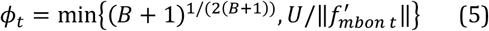

Here,

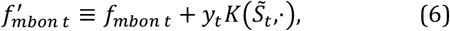

and

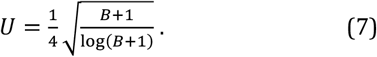

In the original Forgetron, the number of outputs is one. However, in this study, it is extended to multiple outputs. Accordingly, *ϕ*_*t*_ in Eq. 5 is not optimal. To overcome this problem, the value of *ϕ*_*t*_ that minimizes the cumulative error is selected through computational simulations. Therefore, the Model2 fixes *ϕ*_*t*_ to a certain value. This simplification is also suitable as the model for forgetting in biological brain[1][34]. Weight reduction in Forgetron (Model 2) is slightly different from that in Model 1. In Model 1, weight shrinkage is used to detect the age of the oldest kernel. If the weight of the oldest kernel is less than*θ*, it gets removed. In Forgetron, weight shrinkage is used to detect the oldest kernel; however, it is not used to decide whether to remove the oldest kernel. The oldest kernel is removed when the capacity is full. Instead, weight shrinkage reduces the negative impact of removing the oldest kernel.

The model was theoretically analyzed in detail (see **Method**), and the results show that the weight-shrinking mechanism helps reduce the mistake bound of incremental learning on a fixed budget.

Therefore, when Forgetron is used as a model for MBON learning, the forgetting strategy in MBON is essential for learning under limited capacity. The pseudocode is presented in Algorithm 2.

#### Model 3: Least Recently Used

Both Models 1 and 2 prune the oldest kernel when |*S*_*t*_| ≥ *B*. However, age does not always equate to uselessness. Often, the kernels that contribute the least are those used least. If the environment remains stable, the kernels that are accessed least recently are likely to remain rarely used; therefore, it is reasonable to remove the least recently used kernels. To this end, Model3 resets *σ*_*it*_ to its initial value whenever the *i-*th kernel shows the maximum activation among all kernels, provided that the model’s output is correct. Consequently, the the *σ*_*it*_ of the least recently used kernel eventually reaches the minimum value. The pseudocode is presented in Algorithm 3. Note that Model3 employs a spontaneous shrinking mechanism to estimate kernel usage frequency. It reduces weight regardless of output accuracy, using a shrink-ratio *ϕ*^′^such that *ϕ* < *ϕ*^′^ ≤ 1. The model was also theoretically analyzed in detail (see **Method**), and the results showed that mean lifetime (the time interval between the kernel addition and pruning) of each kernel was extended. However, the pruned kernel in the Model3 is not always the oldest one; therefore, Model3’s damage due to pruning is sometimes larger than that of Model2. For this reason, the number of mistakes for the Model3 is not always lower than that of Model2.

### Computer simulation

In this section, we describe the experiments conducted to evaluate model performance in changing environments. These simulations demonstrate that forgetting plays an essential role in reducing cumulative error. The **Method** section presents a theoretical analysis of the proposed models, including the derivation of the crucial shrinkage ratio, which governs their behavior. As the experimental settings differ from theoretical assumptions, the optimal shrinkage ratio is determined empirically. Specifically, since a theoretical proof in such a complex environment has yet to be achieved, we experimentally derive the optimal value for the shrink ratio. First, we outline the assumed environments and their Markov chain formulations. Next, we explain the evaluation procedure used to assess the shrinkage effect in each model. Finally, experimental results are presented.

#### Assumed environments and their Markov chain model

In this section, we describe the assumed environments and their Markov chain models. To predict the sensory inputs from KC outputs, the model must capture the changes in sensory inputs. Sensory inputs are mainly affected by animal actions. Several researchers have investigated the walking patterns of *Drosophila* [13,14] and reported continuous changes in the walking patterns. Accordingly, a model has been developed to generate continuous walking patterns [13], and a Hierarchical Hidden Markov model (HHMM) is used for its analysis [14]. The modeled HHMM shows significantly simpler patterns. Although the model consists of two-layered HMMs, each layer is described by one-dimensional chains of several states. Thus, each state is connected only to its two nearest-neighbor states due to the continuously changing patterns.

Sensory input patterns from KCs are also affected by animal actions. Therefore, sensory input patterns are represented by the HHMM (derivation is provided in the **Hidden Markov model of Kenyon cells’ outputs** section). We use a Markov chain model to generate environmental changes. For simplicity, we assumed a single-layered Markov chain model to evaluate the performance of each model. Therefore, for a five-dimensional generalized sensory input at time *t*, 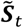 is generated from a multivariate normal distribution whose covariance matrix is *Σ* = *diag*(0.01).

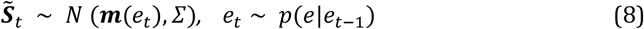

Here, *e*_*t*_ denotes the current state, and *e*_*t*_ ∈ {*e*_1_, *e*_2_, … *e*_40_}. ***m***(*e*_*t*_) denotes the mean vector for the current state and is a fixed random vector defined for each *e*_*i*_ (*i* = 1,2, … ,40). Each cluster center ***m***(*e*_*i*_) is generated from the uniform distribution as follows:

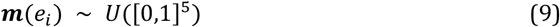

However, if the cluster centers are in proximity to each other, classification becomes difficult. Therefore, we regenerate the cluster centers until the minimum distance between each pair of cluster centers exceeds 0.5. The number of states is set at 40 to ensure that the classification task requires a memory larger than the budget size of each model. State change is defined using a staying state probability *p* function as follows:

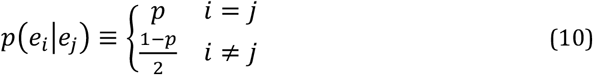

Here, *p*(*e*_*i*_|*e*_*j*_) *i* ≠ *j* is adjusted to fit a real environment, and *p*(*e*_*i*_|*e*_*j*_) = *p*(*e*_*j*_|*e*_*i*_) is set for simplification. This setting is sufficient to evaluate learning performance. Further, we generate nine types of synthetic datasets with *p* = 0.1, 0.2, 0.3,…, 0.9. The size of each dataset is 2000. For each 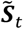, 50 different datasets are prepared, as shown in Fig. 3. In each dataset, each 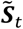 is stored together with its corresponding state vector *y*_*i*_ = *OneHot*(*e*_*i*_), which serves as its label. To fit kernel perceptron learning, the *k*-th element of *y*_*i*_ is 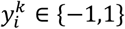. During the computer simulation, the pairs 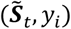 are presented to each model.

**Fig. 3.**
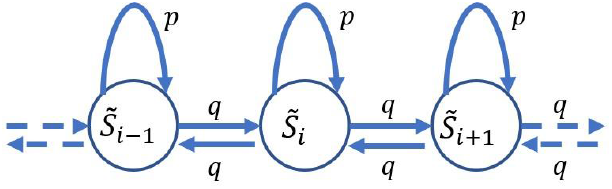
Distributions used in the experiments. Here, *p* denotes the probability of the steady state, *q* denotes the state change probability, and *p* + 2*q* = 1.

#### Evaluation Method

Experiments were repeated at different state-change rates. The cumulative error, which is obtained by summing the differences between the training output and correct labels, was used as an evaluation measure, as in Eq. 11. The experiment was performed to evaluate the MBON model described in a previous study.

**Kernelized incremental learning on a budget as a model of the MBON** *α*’3 **neuron**

For all models, the single-output model was extended to a multiple-output model for the experiments. The multiple outputs used during the simulation were one-hot representations of the predicted labels of the current input vector x(*t*). Therefore, the cumulative error was modified for each round as follows:

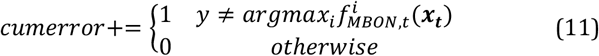

#### Simulation Results

The shared network size, input size, kernel size, and output size were set to 5, 20, and 40, respectively. Various shrinkage ratios were examined to evaluate the effects of weight shrinkage (forgetting)^2^.

##### An example of model behaviors

An example of the cumulative errors of MBON Models 1, 2, and 3 and their competitors (with and without weight shrinkage) at *p* = 0.9 is shown in Fig. 4. The shrinkage ratio *ϕ* was set to 0.64, where the mean cumulative error was approximately minimal for the three models.

**Fig. 4.**
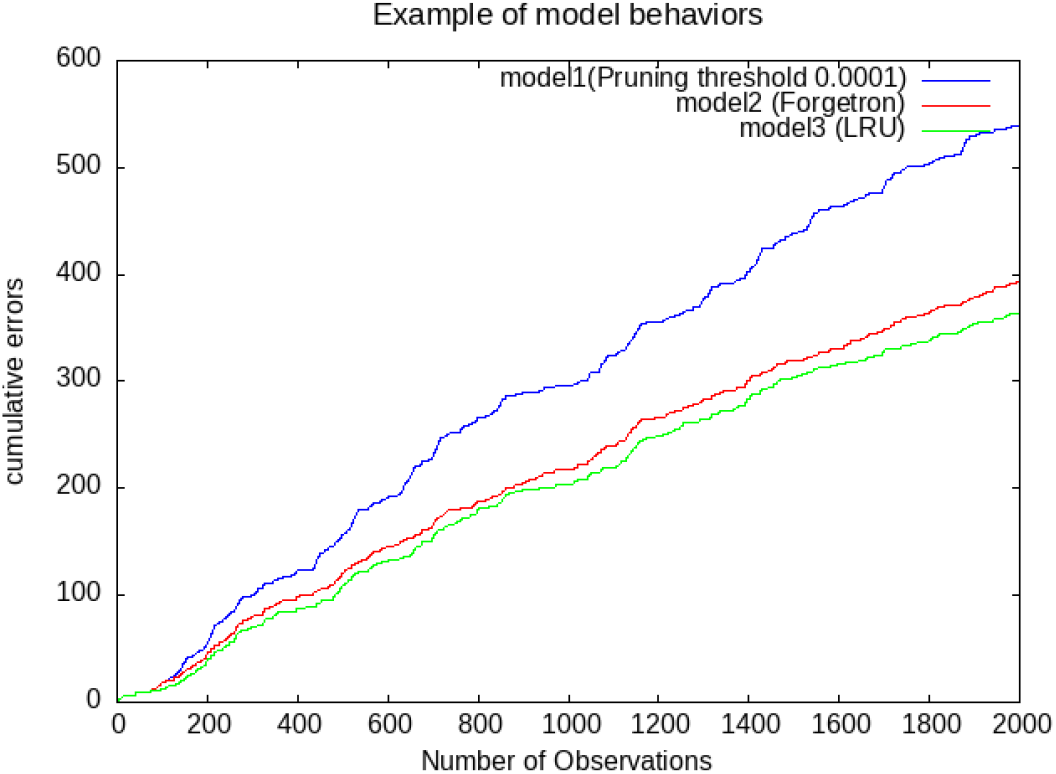
Example of cumulative error vs number of observations (*Budget* = 20, *p* = 0.3) for Models 1, 2 (Forgetron), and 3 (Forgetron with LRU).

During testing, the budget of all models was set to 20, and *γ* was set to 4.0. The parameter used in Algorithm 1 for Model 1 was set as *θ* = 0.1. Therefore, the pruning condition for Model 1 was set as *σ*_*p*_ < 0.0001.

In this example, the cumulative error order was Model 3 *<* Model 2 *<* Model 1. The cumulative error for Model 1 was larger than that for Models 2 and 3. The performance of each model is detailed in the next section.

##### Model 1 sensitivity to weight shrinkage

To evaluate the effect of weight shrinkage (forgetting), the final cumulative errors after learning 2000 samples were calculated for Model 1. Points representing the cumulative error of Model 1 under various weight-shrinkage conditions are shown in Fig 5. The kernel size and budget size were set to 20, and the kernel pruning condition was set at *σ*_*p*_ < *ϵ*, where *ϵ* = 0.01, 0.001, and 0.0001. In Fig. 5, the x-axis represents the shrinkage ratio, while the y-axis represents the mean cumulative error after learning 2000 samples. For each shrinkage-ratio condition, 50 trials were conducted for each of the 90 state-change rates (*p* = 0.1, 0.11, …, 0.99), resulting in 4500 trials. Simulation results (Fig. 5) show a convex curve; therefore, an optimal shrinking coefficient exists at the minimum for each situation, which are around *ϕ* = 0.6 ∼ 0.8. Thus, an appropriate weight shrinkage ratio exists to minimize cumulative errors. Model 1 cannot prune any kernel when *σ*_*p*_ is larger than *ϵ*. Therefore, cases in which Model 1 could not learn the new samples were observed. Therefore, large cumulative errors were observed for a small value of *ϕ* when *ϵ* = 0.0001 and 0.001. The differences between these results are shown in Fig. 7.

**Fig. 5.**
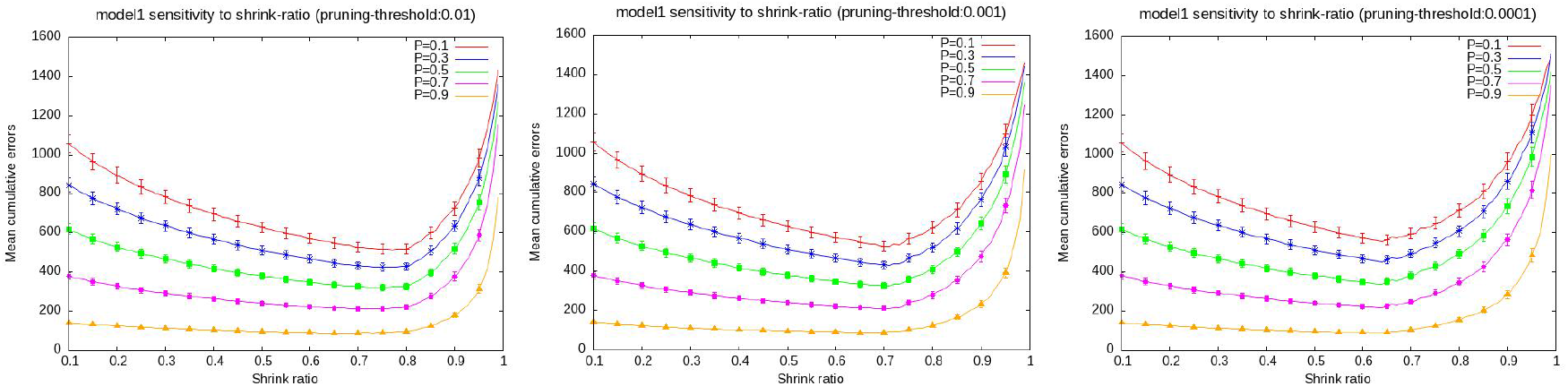
Sensitivity of Model 1 to weight shrinkage after learning 2000 samples. Pruning threshold ranges from *ϵ* = 0.01 to 0.0001.

##### Model 2 (Forgetron) sensitivity to weight shrinkage

Further, we evaluated the effect of weight shrinkage (forgetting) on Model 2 (Forgetron). Similar to the previous section, the cumulative errors after learning 2000 samples were analyzed. The cumulative error for each shrinkage ratio is shown in Fig. 6. Under all conditions, the cumulative errors were minimal for *ϕ* = 0.64–0.77.

**Fig. 6.**
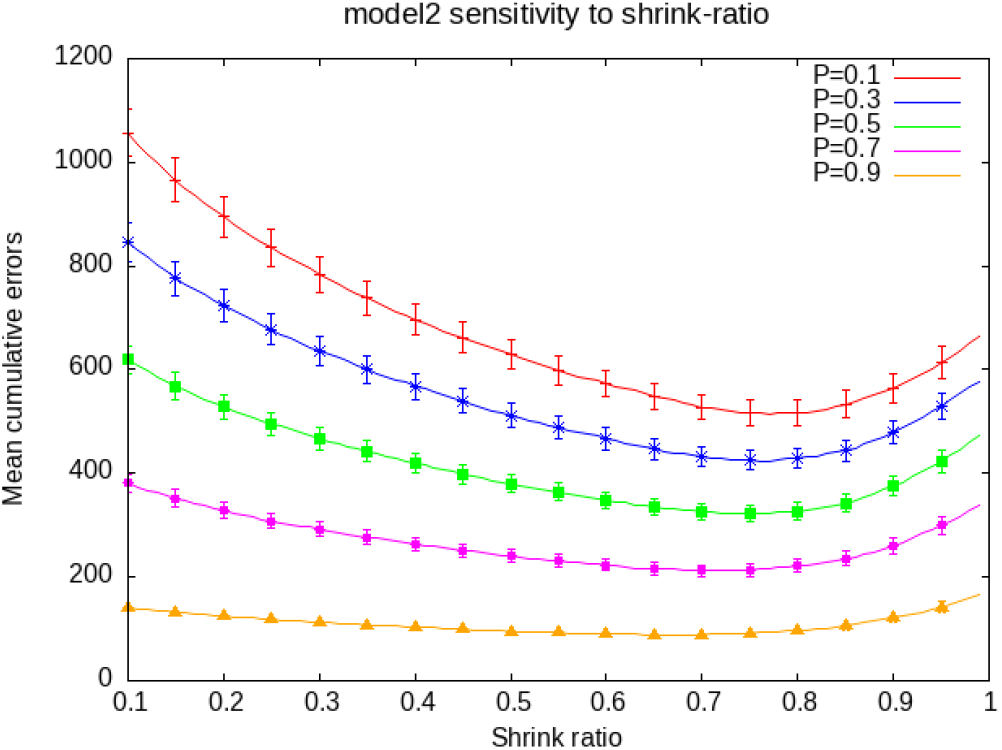
Model 2 (Forgetron) sensitivity to weight shrinkage after the learning of 2000 samples. State stay rate: *p* = 0.1 to *p* = 0.9. The errorbars denote 95% confidence intervals.

###### Model 1 vs Model 2 (Forgetron)

We compared the performance of Models 1 and 2 (Forgetron). Similar to the previous sections, the cumulative errors after learning 2000 samples were analyzed. The cumulative error of Models 1 and 2 with pruning threshold *ϵ* = 0.05, 0.001, and 0.0001 is shown in Fig. 7.

**Fig. 7.**
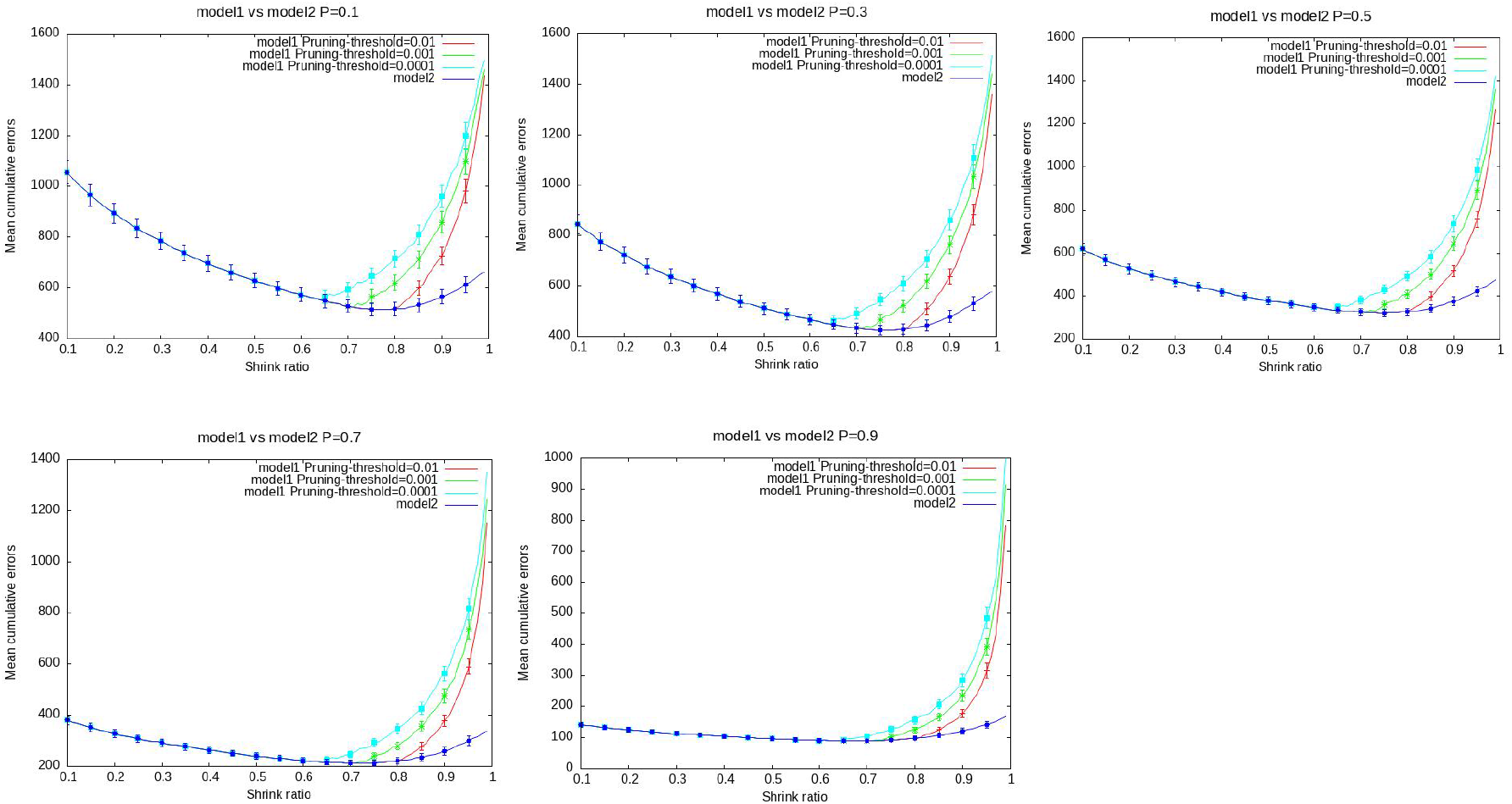
Comparison of cumulative errors between Models 1 and 2 after learning 2000 samples. The pruning threshold of Model 1 is *ϵ* = 0.01, 0.001, and 0.0001. The errorbars denote 95% confidence intervals.

The cumulative error of Model 2 was partly identical to that of Model 1. However, the cumulative errors of Model 1 were larger than those of Model 2 in most cases where the pruning threshold *ϵ* was small. Thus, Model 1 cannot prune any kernels when *σ*_*p*_ is consistently larger than *ϵ*.

###### Model 3: LRU vs Model2 (Forgetron)

We conducted a comparative study between Model 2 (Forgetron) and Model 3 (Forgetron with LRU). The cumulative errors after learning 2000 samples were analyzed by sweeping the shrink ratio in the interval [0.7,0.8],where the cumulative error of Model2 is nearly minimum. The cumulative error points for (*Model* 2, *Model* 3) are shown in Fig. 8. The points below the line *y* = *x* indicated that the cumulative errors of Model 3 were smaller than those of Model 2. The distribution of points was slightly more concentrated on the lower side. Specifically, the proportion of points below *y* = *x* was varied depending on the spontaneous shrink ratio *ϕ*^′^. For example, in the case of *ϕ*^′^ = 0.99, 82.7% of the 550 × 9 trials (9 variations in stay probability: 0.1, 0.2 …0.9). But the ratio was reduced when *ϕ*^′^is closed to 1(see Table1). Thus, the LRU strategy performed relatively better than Model 2 (Forgetron). However, the LRU model predicted the future usage of each memory based on its past usage history. The prediction was sometimes incorrect owing to variations in the inputs. In such cases, the cumulative error in Model 3 increased. Moreover, the removed kernel was not necessarily the oldest kernel; therefore, the impact of its removal was greater than that in Model 2.

**Fig. 8.**
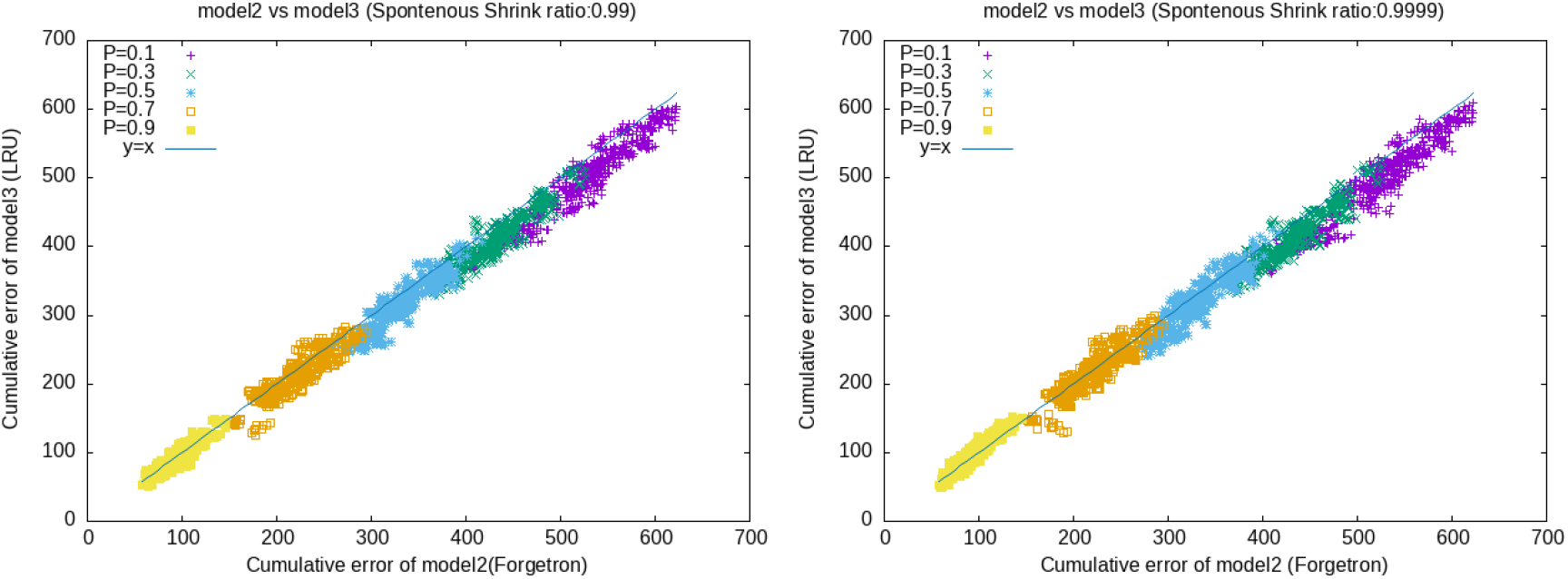
Comparison of cumulative errors between model2 (Forgetron) and model3 (Forgetron with LRU) after learning 2000 samples. Shrink ratio was sweeped from 0.7 to 0.8, where the cumulative error of model2 is nearly the smallest. The spontaneous shrinkratio was *ϕ*^′^ = 0.99 (left) and 0.9999(right). For *P* ∈ {0.9, 0.7, 0.5, 0.3, 0.1}, the proportions of points below *y* = *x* across 550 trials were 57.2%, 76.7%, 84.9%, 92.7% and 93.8% for the left case, whereas these proportions were 36%,58.5%,73.6%,90.7% and 94.5% in the right case, respectively.

**Table.1.** Proportion of Model 3 superiority vs. spontaneous shrink ratio.

| $\phi'$ | P=0.1 | P=0.2 | P=0.3 | P=0.4 | P=0.5 | P=0.6 | P=0.7 | P=0.8 | P=0.9 | average |
| --- | --- | --- | --- | --- | --- | --- | --- | --- | --- | --- |
| $\phi'=0.99$ | 0.94 | 0.96 | 0.93 | 0.96 | 0.85 | 0.80 | 0.77 | 0.66 | 0.57 | 0.83 |
| $\phi'=0.999$ | 0.95 | 0.97 | 0.91 | 0.92 | 0.75 | 0.69 | 0.61 | 0.50 | 0.41 | 0.75 |
| $\phi'=0.9999$ | 0.95 | 0.95 | 0.91 | 0.90 | 0.74 | 0.68 | 0.59 | 0.46 | 0.36 | 0.73 |

**Table 2:** Comparison of number of mistakes for the LRU and Forgetron models.

| $e_i$ | Average Number of mistakes of both models | Pruning damage | Mistakes under the LRU strategy VS under the Forgetron strategy |
| --- | --- | --- | --- |
| $e_i \rightarrow 1$ | Small | LRU>Forgetron | LRU ≈ Forgetron |
| $e_i \ll 1$ | Large | LRU<Forgetron | LRU < Forgetron |

This interpretation corresponds to the results shown Fig. 8.

## Discussion

### Relationships with other approaches

We constructed the kernelized MBON*α*^′^3model to explain its need for forgetting in novelty detection and learning. This model is not related to reward signals. However, almost all previous MBON models are related to reward signals [15,16]. A new model [17] developed to predict olfactory learning in *Drosophila* was used to predict the aversive or appetitive valences of odors. The model did not follow the free energy principle proposed previously by Friston et al. [18]. Instead, the design principle was based on reducing unfamiliar situations.

In existing literature [16,17], two models have been designed based on the behavior of MBON*α*’3 and novel detection mechanisms. Researchers have developed models based on the prediction of agreeable stimuli [17]. Such models do not follow the free-energy principle [18]. Instead, the design principle is based on reducing surprise. Therefore, such models were designed to fit empirical data.

In this study, the model was designed using a learning algorithm with a fixed budget. The proposed model was used to analyze the error bound under a fixed budget. Therefore, the results did not directly address the two models [16,17] but instead presented new insights into MBON models from the perspective of minimizing cumulative errors under a fixed budget.

Three forgetting models were considered for this study. However, another possibility was emphasized. Forgetting redundant memory serves as a second possible approach. Existing MBON models [15,16] are based on reward signals. A general model was proposed for MBON *α*’3 and other areas. From the perspective of behavior, the activity of MBON*α*’3 was not directly related to reward signals.

### Biological relevance of the three models

Three candidate MBON models with different kernel-pruning strategies have been discussed. However, additional behavioral experiments are required to determine the most appropriate model. Based on the model error counts, Models 2 and 3 perform excellently. This section discusses the application of the obtained results to the *Drosophila* brain.

#### Model 1 vs Model 2

To compare the performances of Models 1 and 2, an overloaded learning task was designed, in which more than 20 odors were sequentially presented in our simulations.

Theoretically, if MBON learning in *Drosophila* follows Model 1, the system would ignore novel odors after presenting approximately 10–20 odors without rest. Conversely, if the learning follows Model 2, *Drosophila* should continue to respond to novel odors even after extensive learning. From the perspective of the free energy principle [18], Model 2 served as a more plausible approach than Model 1.

Our simulation results demonstrated that Model 1 terminated learning when novel odors were presented, leading to higher prediction errors. In contrast, Model 2 maintained a larger set of learned odors, which allowed the system to better predict incoming sensory inputs. The experimental results shown in Fig 7 also clearly suggest that the cumulative error of Model 2 is significantly lower than that of Model 1. In summary, while these findings demonstrate the superior predictive performance of Model 2, further behavioral studies are required to confirm whether actual *Drosophila* learning aligns with the dynamics of Model 2.

#### Model 3 vs Model 2

Model 3 incorporates the LRU pruning strategy. In our simulations, a subset of previously learned odors was presented intermittently during a novel-odor learning task. To discuss which model is more appropriate, we must consider actual *Drosophila* behavior. For instance, when a fly shows alerting behaviors to a frequently presented old odor, this pattern is consistent with Model 2. Conversely, if the fly does not show such behaviors, its learning potentially follows Model 3. Experimental results in Fig 8 suggest that Model 3 outperforms Model 2 when the state-staying probability is small. While Model 3 showed superior average performance in our simulations, the biological validity of these models should be evaluated based on future observations of real *Drosophila* behaviors.

### Comparison between the results of theoretical analysis and computer simulations

Our sensitivity analysis demonstrates that the model effectively manages to minimize the cumulative errors when an appropriate shrink ratio is applied. Importantly, these empirical results are consistent with our theoretical findings, which guarantee the existence of a unique optimal shrink ratio *ϕ*_*_ within the interval (0, 1).

### Validity of learning without the reward signal

MBON*α*’3 neurons are activated by novel odor stimuli. Although existing computational models for MBONs are usually based on reinforcement learning, they are unsuitable for MBON*α*’3 neurons. Since a novel stimulus cannot be guaranteed to be associated with a reward, an MBON model is proposed without explicit reward signals.

We assume that MBON*α*’3 behaviors can be determined following the free-energy principle proposed by Friston et al. [18,19]. For instance, Isomura [20] adopted the free-energy method and provided a statistical interpretation. The free-energy principle is used to model biological behavior under the biologically inferred assumption that actions and learning are performed to reduce surprises or unexpected situations. Interestingly, free-energy-principle-based computational models do not always assume the presence of reward signals.

In the free-energy method, biologically inferred learning systems are assumed to maximize.

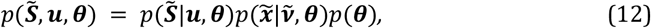

where 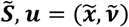, and *θ* denote a generalized sensory input vector, generalized hidden state vector, and parameter vector, respectively^3^.

Although the system is assumed to minimize surprises, defined as 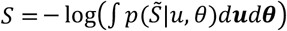, the direct minimization of *S* is difficult to execute. Practically, surprises can be reduced by minimizing the free-energy function [21].

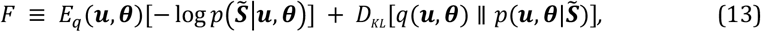

where *D*_*KL*_ () is the Kullback–Leibler divergence, *q*(*u, θ*) denotes a recognition distribution, and 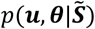 denotes a true u distribution.

Under the free-energy principle, we assume the existence of a generative model that represents 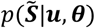. MBONs exhibit backward connections to the antennal lobe (AL) [22]. The backward connections can be represented in the generative model 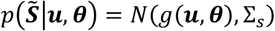 of the *Drosophila* brain (Fig. 9).

**Fig. 9.**
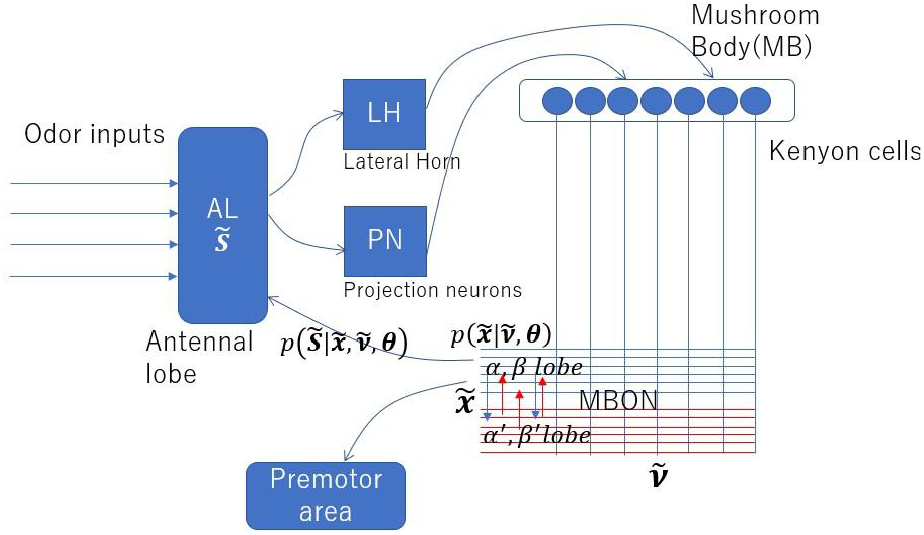
Connectivity of mushroom body output neurons and the free-energy principle. (Extended figure in [23]).

The MBON*α*’3 neuron is assumed to be a part of the generative model for the hidden-state vector 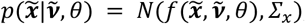, where the cause 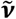 is assumed to be the output vector from KCs.

### Hebbian learning minimizes free-energy

We assume that 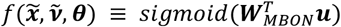, where ***W***_***MBON***_ denotes the weight ***W*** vector of the MBON *α*’3 neuron. The change in ***W***_***MBON***_ can be denoted as a Hebbian learning function, described as [21,24]

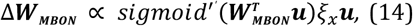

where *ξ*_*x*_ denotes the output from an error-coding neuron [18,21]^4^. Although we assume that the model is a single-layer network, actual MBONs exhibit multi-layered structures [1]. Thus, the model can be improved to a multi-layered version as [24].

According to existing literature [2], MBON*α*’3 outputs are adjusted by anti-Hebbian learning, which is triggered by the activation of *α*’3 dopaminergic neurons. Assuming that anti-Hebbian learning can be reformulated as Hebbian learning, and that activation of *α*’3 dopaminergic neurons represents *ξ*_*x*_ in Eq. 14, then MBON*α*’3 neurons can acquire learned outputs from KCs to reduce free energy.

If the learning model does not recognize the current stimulus, the surprise term increases. Consequently, the model attempts to learn to recognize the current stimulus and restore recognition. Thus, the system should encode the current stimulus.

## Methods

In Eqs. 3 and 15, *σ*_*j,t*_ denotes a decay term such that *σ*_*jt*_ ≤ 1. Generally, *σ*_*jt*_is assumed to decrease at each round at a certain ratio, thereby gradually reducing the output value of the MBON model. The old kernel exhibits the smallest *σ*_*jt*_. Thus, we can remove the oldest kernel by identifying the kernel with the smallest *σ*_*it*_ value. However, the same result can be achieved by simply removing the oldest kernel without shrinking the weights. The differences between the two methods—removing the oldest kernel with and without forgetting—were analyzed to clarify the effects of shrinkage (forgetting).

### Simplification of the proposed model

To theoretically assess the incremental learning capacity under a budget constraint, we adopted the theoretical analysis of the Forgetron proposed by Dekel et al. [3]. Accordingly, we simplified the developed model without losing generality. The multidimensional output of the model was switched to one-dimensional output.

The learning samples consisted of input vectors ***x***_***t***_ ∈ *R*^*n*^ and the corresponding label vectors *y*_*t*_ ∈ {−1, +1}. When the model predicted the correct label, the current input vector ***x***_***t***_ was not regarded as a novel input (a new kernel was not allocated). Thus, the model was simplified to a kernel perceptron.

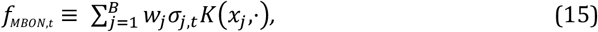

where *K*(*x*_*j*_,·) is an infinite-dimensional vector converted from *x*_*j*_, and *B* denotes the upper bound of the number of kernels. Notably, *f*_*MBON*_(*x*) = ⟨*f*_*MBON*_, *K*(*x*,·)⟩, where ⟨·,·⟩ denotes dot product operation. Similar to the proposed model, *f*_*MBON*_ learns samples *X* = [*x*_*t*_, *y*_*t*_]^*T*^ online. In this analysis, *f*_*MBoN*_ was evaluated using a *hinge-loss* function.

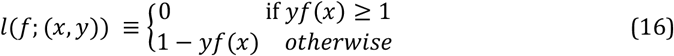

The kernel perceptron alternately executed inference and learning, as shown in Fig. 2. The learning was performed to minimize the cumulative error (number of mistakes).

#### Mistake bound of removing the oldest kernel with forgetting

We considered the mistake bound of the kernel perceptron by removing the oldest kernel while forgetting. Dekel et al. [3] demonstrated the mistake-bound *M* in Theorem 6.2 in their manuscript. However, Theorem 6.2 should be satisfied under the condition *B* ≥ 83, which is much larger than the estimated capacity of MBON:20. Therefore, the more general mistake bound presented in Lemma 5.2 was used [3]. Moreover, the original Forgetron uses adaptive shrink ratio. In this study, we assume that the shrink ratio is fixed to align with the biological simplicity of dopaminergic forgetting mechanisms in the Drosophila brain [1][34]. In our previous study, we used this lemma to analyze the mistake bounds of the MBON model [4]. As the analysis in the previous study lacked rigor, we reanalyzed the model in this study. The following assumptions were made:

- Let *g* be a desired competitor function in the Hilbert space *H*_*k*_ such that |*g*| ≤ *U*, where *U* is defined in Eq. 7.
- Let *ϕ*_*t*_ ≤ 1 be the shrinking coefficient for *σ*_*i,t*_.
- Let (*x*_1_, *y*_1_),…, (*x*_*T*_, *y*_*T*_) be a sequence of examples such that *K*(*x*_*t*_, *x*_*t*_) ≤ 1 for all *t*.
- Let *J* be a set of examples in which *f*_*MBON*_ mistakes the predicted label *J* = {*t*|*y*_*t*_ *f*_*MBON*_ (*x*_*t*_) < 0}.
- Let 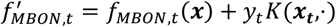
- Let 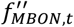 be 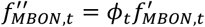.
- Ψ_*t*_ be the magnitude of loss due to pruning the old kernel and

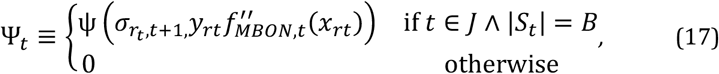

where *Ψ* (*λ*, μ) ≡ *λ*^2^ + 2*λ* − 2*λ*µ, and the suffix *r*_*t*_ denotes the removed kernel at *t*. Then, the number of mistakes of *f*_*MBON*_ satisfies the following inequality.

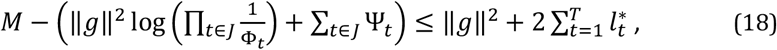

where *g* denotes the desired competitor function; 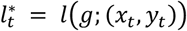. A detailed derivation of Eq. 18 is presented in a previous study [3]. From the results, *M* is larger than the mistake bound of the original kernel perceptron, which does not use shrinking or pruning kernels: 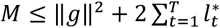. In basic Forgetron (described in Fig. 5.1 of [3]), *ϕ*_*t*_ is adaptively controlled such that *ϕ*_*t*_ is variable. By adapting *ϕ*_*t*_, Dekel et al. [3] showed that the number of prediction mistakes of the basic Forgetron is bounded as 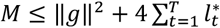. To investigate the real duty of forgetting in the *Drosophila* MBON, we assumed that the value of *ϕ*_*t*_be fixed to *ϕ* ≤ 1 for all *t*. To evaluate the mistake bound with a constant shrink ratio *ϕ*, we start by estimating the deviation from the adaptive Φ_*t*_ used in Dekel et al. [3] This estimation, as shown in the following Lemma 1, provides the basis for our subsequent proof.

##### Lemma 1.

*Using a shrink ratio ϕ* < 1, Φ_*t*_*can be defined as*

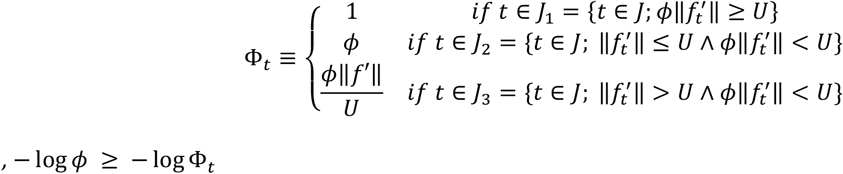

*Proof*. Since *ϕ* < 1, log Φ_*t*_ *=* 0 where *t* ∈ *J*_1_; log Φ_*t*_ = log *ϕ* when *t* ∈ *J*_2_; 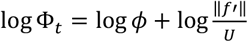 when *t* ∈ *J*_3_. log Φ_*t*_ ≥ log *ϕ*. Thus, log *ϕ* ≥ log − Φ_*t*_

Notably, the MBON model reduces the mistake bound when *ϕ* < 1, and shrinking (forgetting) is essential to maintain the mistake bound low.

By using ∥ *g* ∥≤ *U* and |*J*| ≤ *T*, we derive the lemma.

##### Lemma 2.

*ϕ denotes the shrinking coefficient for the decay ratio of σ*_*i,t*_, ∀*i* ∈ *S*_*t*_, *and* 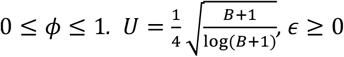 *The number of mistakes M is bounded by*

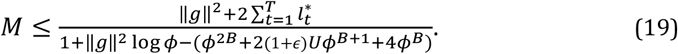

*Proof*. Since 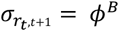, Ψ can be rewritten as follows:

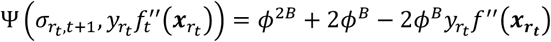

From the Cauchy–Schwarz inequality, 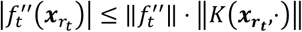 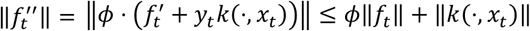. Under a fixed shrink ratio *ϕ*, there are possibilities that 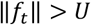. Thus, 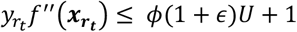 when 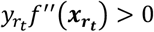 and 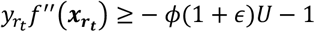 when 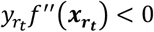, where *ϵ* ≥ 0. Therefore, we can rewrite the expression as follows:

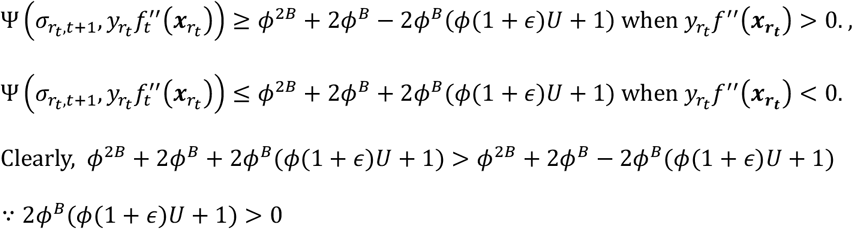

After all, 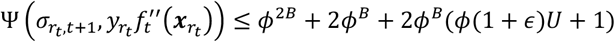

From Lemma 1, we rewrite Eq. 18 as follows:

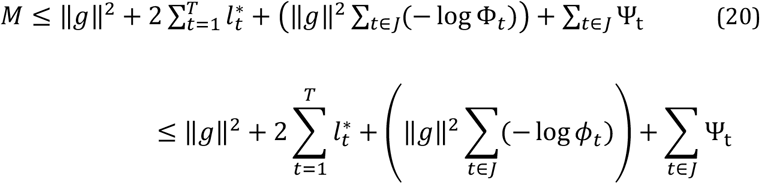

where the right-hand side is also bounded by

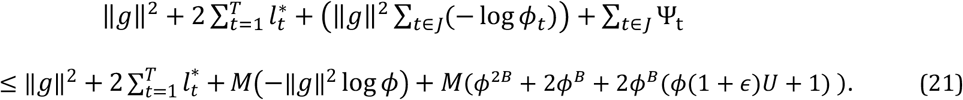

Here, Eqs. 20 and 21 yield

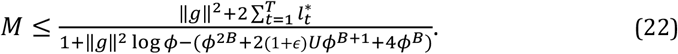

The denominator of the right-hand side of Eq. 22 depends on *ϕ*. Thus, we can optimize *ϕ* to minimize the mistake bound. The upper bound of the number of mistakes is minimized when *ϕ* is set to a certain value, as shown in Theorem 1. An example of the relationship between *ϕ* and the mistake bound is shown in Fig 10. From Lemma 2, we obtain Theorem 1.

**Fig. 10.**
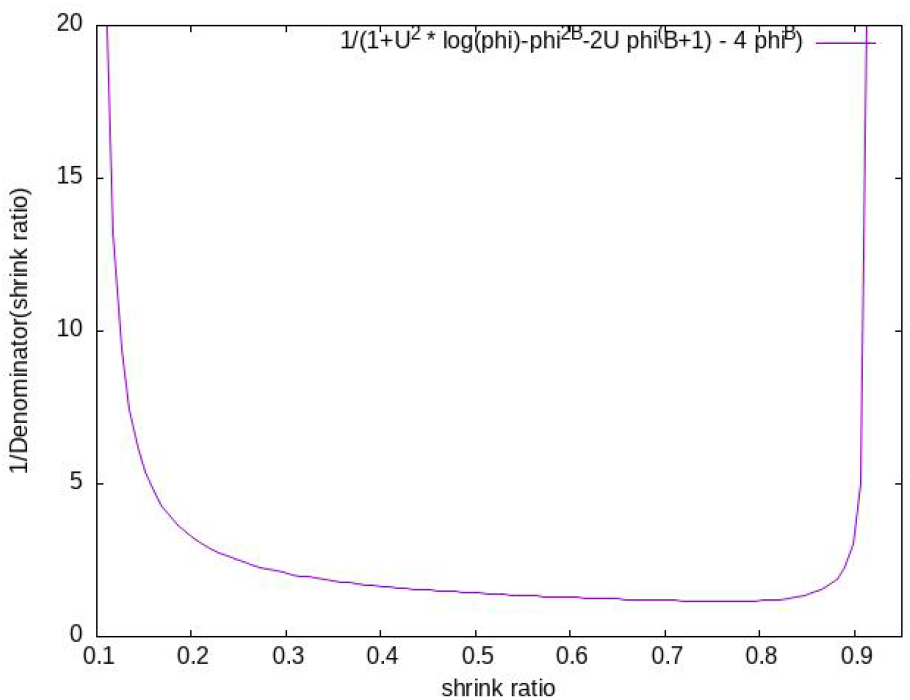
Example of a rough relationship between the shrinking coefficient *ϕ* and right-hand side of Eq. 22. The numerator was set with 1 and ‖*g*‖^2^with *U* to visualize the relationship. The mistake bound is minimized when *ϕ* = *ϕ*_*_ < 1. In this example, *B* = 20 and 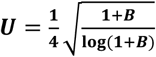.

##### Theorem 1.

Let *f*_*MBON*_ be a kernel perceptron with a budget *B* and a fixed shrinking coefficient *ϕ* for the decay of *σ* (∀*i* ∈ *S*_*t*_). Assuming ; *T* > *B, and* 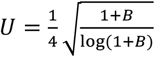, there exists an optimal shrinking coefficient *ϕ*_*_ in the interval 0 < *ϕ*_*_ < 1 that minimizes the upper bound of the number of mistakes made by *f*_*MBON*_.

*Proof*. Let *D*(*ϕ*) be the denominator of the right-hand side of Eq. 22.

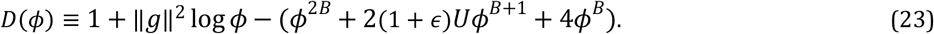

The first and second derivatives of *D*(*ϕ*) with respect to *ϕ* are obtained as follows:

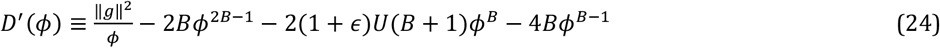

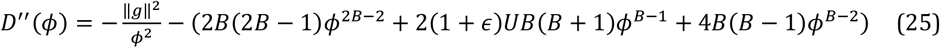

Since all parameters ‖*g*‖^2^, *B,U*, and c are positive and *B* > 1, it is evident that *D*^′′^(*ϕ*) < 0 for all *ϕ* ∈ (0,1]. This implies that *D*(*ϕ*)is strictly convex on this interval.

Regarding the boundary behavior of the first derivative, Eq. 24 indicates that *D*^′^(*ϕ*) → ∞ as *ϕ* → 0^+^.

Conversely, as *ϕ* → 1^−^, we have

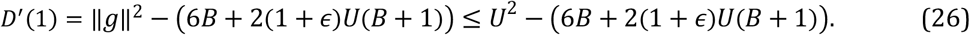

Given that 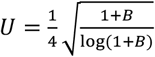, it follows that *U*^2^ < *B*, for *B* > 1, ensuring *D*^′^(1) < 0. By the intermediate value theorem and the strict convex of *D*(*ϕ*), there exists a unique optimal *ϕ*_*_ ∈ (0,1) that minimizes the mistake bound.

This theorem indicates that the MBON model must forget memories at a certain ratio to reduce the number of prediction mistakes.

### Analysis of the MBON model with the least recently used strategy for the pruning hypothesis

The LRU strategy is a well-known page replacement algorithm in computer science. In this section, we analyze the performance of the LRU strategy in terms of the average lifetime. In the analogy between page replacement algorithms and kernel-based online learning algorithms, each page represents a kernel, and the memory budget represents the maximum number of kernels that can be stored in memory. Consider a string reference call *w* = {*r*_1_, *r*_2_,… } which describes the order of call of the pages of an algorithm *A*.

This string reference call corresponds to the order of appearance of the classes of input vectors in the online learning algorithm. The input vector ***x***_***i***_ is generated from the *i* th state, where we assume that *i* = *argmax*_*j*_ ***K(x***_***i***_, ***x***_***j***_) represents each *r*_*i*_ as a kernel. There are *k* kernels, and cardinality |*w*| = *n*. It is given that *A* works under the budget *B*.

Converting each algorithm A to its irreducible Markov Chain with a fixed stationary state is crucial to calculate its mistake bound relative to its performance. We use the ergodic theorem.

Let f(·) be a bounded function from *X* → ℝ, with *X* being a superset of *x*_*i*_ and ℝ being the set of all real numbers. If (*x*_0_, *x*_1_,…, *x*_*n*_) is an irreducible, time-homogeneous, and discrete Markov Chain, then

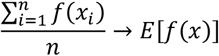

as *n* → ∞, almost surely with probability 1. *Sπ* = *π*, where *S* be the transition matrix of this Markov Chain, and π = [e_1_, e_2_,…, e_n_]^T^ is the stationary state of the Markov Chain. The element *e*_*i*_ represents the ergodic numbers (or probabilities) as *t* → ∞, and it is stable (which justifies the existence of *d*_*avg*_, later on). *n* is assumed to be a large number.

*P*_*ij*_ denote the state transition probability from state *i* to state*j*

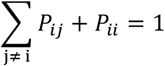

Let *x* = ∑_*i*≠*j*_ *P*_*ij*_ *e*_*j*_, and *y* = ∑_*i*_ *P*_*ii*_ *e*_*i*_. On closer inspection, we can see that *x* is the probability of the switching between states representing its “volatility” relative to its environment, and *y* is the probability of the fly remaining in the same state as earlier representing its “stay” relative to its environment.

Let us now view the specific Markov Chain in terms of reference calls. Let *D* = (*d*_0_, *d*_1_,…, *d*_*n*_) be the increase in lifetime of each kernel with recovery that occurs at every *O*(*ω*) = 0 for a string reference call *ω*, where *O*(*ω*) is a function from *ω* → *O*, where elements *o* of *O* are {0, 1}, and *O*(*ω*) converts each string *ω* into binary by individually labeling all first and unique instances of a kernel as 1, else 0 if it has been called in the past. We define |*D*| = *Z*, where *Z* is the number of zeros in the *O*(*ω*) operation and |·| represents the cardinality of the set *D*.

It is observed that kernel recoveries occur in the 0 s of this binary *O*(*ω*). After each step of the algorithm, the pages (kernels) arrange themselves in the order of their priority, and the increments to page *i* are represented as *d*_*i*_.

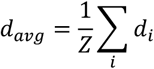

If *T* samples are presented and *i*th pattern appears in the reference call *ω*, it appears *T*· *e*_*i*_ times under budget *B*. The approximate window of *i* th pattern recurrence in string *ω* is *T/e*_*i*_, for *A* that has no definite recoveries in the lifetime of pages (kernels).

The *i* th pattern recurrence in string *ω* is *d*_*avg*_ + *T*· *e*_*i*_ for *A* that has definite recoveries in the lifetime of pages (kernels), averaging as *d*_*avg*_ over the string. This will be our indicator of performance, showing the degree of performance convergence between LRU and Forgetron.

An algorithm with conscious and consistent recoveries in pages (kernels) will have *d*_*avg*_ > 0. For Forgetron, *d*_*avg*_ = 0; hence, the average lifetime of a kernel (page *i*) is longer for LRU than Forgetron. This means that LRU’s mistakes caused by a kernel pruning is less than that of Forgetron; hence, the LRU model is expected to perform better if the damage caused by pruning and weight shrinking is not too high.

Analogous to cache theory, on each instance of *y*, the LRU resets its corresponding weight to 1; it can also be observed as the probability of “resetting” of memory, for every next page reference. Similarly, *x* is the probability of “decaying” of memory, for every next page reference.

Notably, when *e*_*i*_ is low, it indicates a natural low recurrence environment, where the states are not clustered or periodic but rather chaotic and random. Here, our indicator tends to a lower value (indicating “non-convergence” in performance), and LRU is expected to perform better than Forgetron, given *d*_*avg*_ + *T* · *e*_*i*_ > 0 for LRU and *d*_*avg*_ = 0 for Forgetron.

Furthermore, a low “stay” probability (y) leads to performance degradation for both algorithms due to the random nature of the inputs. However, the performance disparity becomes more pronounced, with the probability of LRU outperforming Forgetron (P (LRU > Forgetron)) exceeding 0.9. (see purple, green points in Fig. 8). This phenomenon suggests that Forgetron’s error-driven update mechanism may be particularly sensitive to input volatility—a point that warrants further theoretical or empirical investigation.

A notable property of the Forgetron algorithm is that weight shrinkage occurs exclusively upon incorrect classification. Consequently, a high “stay” probability (y) naturally enhances its accuracy owing to the repeated presentation of similar inputs, which minimizes the performance gap where the LRU model might otherwise outperform Forgetron (P (LRU >Forgetron) is limited to 0.57 for the spontaneous shrinkratio *ϕ*^′^ = 0.99 in our observation) (see yellow points in Fig.8 and Table1.).

#### Damages caused by weight shrinking for LRU

Notably, weight shrinking is applied not only when the classification is correct but also when it is incorrect, because this shrinkage is essential to realize the least recently used strategy. This means that the damage caused by weight shrinkage is larger than that of the original Forgetron. To reduce the damage caused by weight shrinking during correct classifications, the weight shrinking ratio for the correct classification should be larger than that for the incorrect classification. By this modification, we can ignore the damage caused by weight shrinking during the correct classification.

For the insect brain model, this modification is reasonable because the dopamine neurons in the insect brain are activated only when the predicted label is incorrect.

Moreover, the data in [2] also suggest that the MBON*α*’3 neurons forget all odor memories for about 1 hour after the conditioning.

This means that forgetting without any stimulation in the insect brain really occurred, corresponding to the weight shrinking during correct classification.

However, the LRU strategy still causes larger damage during incorrect classification because the LRU kernel is not always the oldest kernel.

To analyze this effect, we define 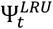 as the loss due to the pruning of the LRU kernel at 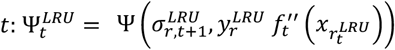, where 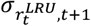 denotes the decay term of the LRU kernel at *t*.

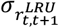 is larger than 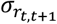 when the LRU kernel is not the oldest kernel, whereas 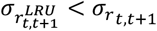 when the LRU kernel is not activated more than *B* times.

Thus, the damage caused by pruning of the LRU kernel is validated depending on the situation.

In the worst case, the LRU strategy repeats correct classifications to activate all kernels uniformly and then misclassifies the input that activates the LRU kernel. In such a case, the damage caused by pruning the LRU kernel is nearly equal to 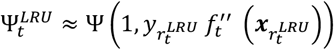.

Note that, in the LRU strategy, the pruned kernel’s 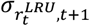 is at least 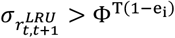. Conversely, in the case of Forgetron, the pruning target parameter 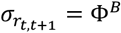. Therefore, if 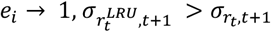. Thus, the difference depends on *e*_*i*_.

If *e*_*i*_ is large, the pruning damage is correspondingly greater. We ca argue that in more “natural” environments, patterns are less, and randomness is higher. In such settings, a lower “stay” probability is more appropriate, suggesting that the LRU is superior to the Forgetron under “natural” conditions. This interpretation is consistent with our simulations.

#### Tradeoff between the pruning damage and LRU effect

Thus, the number of mistakes of the LRU is greatly valid for the situation, and the mistake bound is hardly analyzed generally. Summarizing the discussion, we obtain the following table.

#### Spontaneous shrinkage reduces the pruning damage

Note that the discussion so far has not accounted for the spontaneous shrink ratio *ϕ*^′^, where *ϕ* < *ϕ*^′^ < 1. This ratio *ϕ*^′^ serves to monitor kernel usage intensity even when Model3 (LRU) yields a correct classification.

Interestingly, *ϕ*^′^ also helps mitigate the damage caused by pruning a kernel. Recall the loss incurred by pruning of the LRU kernel at time *t* is 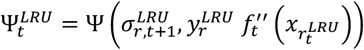. The magnitude of 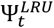 depends on 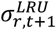; specifically, a smaller 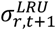 results in a reduced 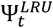. Since all kernel’s 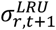 are multiplied by *ϕ*^′^, a smaller *ϕ*^′^faciliates the reduction of 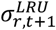 even during correct classifications, thereby minimizing 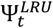 as shown in Fig.8 and Table1 results. By leveraging spontaneous weight shrinkage, we can surgically remove target kernels while shielding essential memories from unintended decay.

## Supporting information

Additional derivations and equations supporting the hypotheses in the manuscript.

## Code Availability

The source code for the models and simulations presented in this study is publicly available on GitHub at [URL: https://github.com/sus8818-beep/MBONA3models] (<u>DOI: 10.5281/zenodo.19571114</u>). The repository includes all scripts necessary to reproduce the results and figures.

## Acknowledgements

*We would like to thank Editage (www.editage.jp) for English language editing*.

## Author contributions

Conceptualization: Koichiro Yamauchi

Data curation: Koichiro Yamauchi

Formal Analysis for Models 1 and 2: Koichiro Yamauchi

Formal Analysis for Model 3: Aditya Gyanprakash Nirmale

Software: Koichiro Yamauchi

Writing - original draft: Koichiro Yamauchi

Writing - review & editing: Koichiro Yamauchi, Aditya Gyanprakash Nirmale

Funding acquisition: Koichiro Yamauchi

Supervision: Koichiro Yamauchi

---

Fig.S1: An example of before:(a) and after:(b) snapshots of applying Eq 39 and Eq 40 in the case of Hebbian learning. The red colored kenyon cells are activated cells and they represents the spase feature vector of presented odor. Such weight changes are triggered by the activation of the DANs.

## Footnotes

1 The support set is the set of kernels.

2 From Theorem 1 and its proof, we determine *ϕ* using Eq 23. However, in this simulation setting, the MBON model has 40 outputs, which does not directly correspond to the Theorem. Therefore, we examine various values of *ϕ* in the simulation.

3 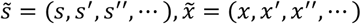 and 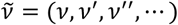

4 Strictly speaking, we have to consider the integral of 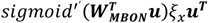 from *t* = 0 to *t* to derivate the change in the parameters. Moreover, the gradient of a prior distribution *p*(W_*MBON*_) is also should be included. However, these operators are omitted for simplicity.

