## Additional derivations and equations supporting the hypotheses in the manuscript. for "Learning by forgetting: A computational model of insect brain"

### Supporting information

#### Hidden Markov model of Kenyon cells' outputs

Let us assume that Drosophila behavior can be represented by a Hidden Markov model as suggested by Tao et al. [14]. Let us denote the Drosophila's current state as $\mathrm{x}(t)$, which is determined by its previous state and actions.

$$\boldsymbol{x}(t) \sim p(\boldsymbol{x}|\boldsymbol{x}(t-1), \boldsymbol{A}(t-1)) \tag{26}$$

The action of Drosophila at time $t$, $\boldsymbol{A}(t)$, is also influenced by the Kenyon cell output $\boldsymbol{k}(t)$.

$$\boldsymbol{A}(t) \sim p(\boldsymbol{A}|\boldsymbol{A}(t-1), \boldsymbol{k}(t-1)) \tag{27}$$

Tao et al. [14] showed that this transition can be represented using a Hidden Markov model. The state $\mathrm{x}(t)$ is observed through the animal's sensors, and Kenyon cells represent it as

$$\boldsymbol{k}(t) = Kenyon(\boldsymbol{x}(t)), \tag{28}$$

where $Kenyon(\boldsymbol{x}(t))$ denotes the Kenyon cell representation of the current state. The responses from the Kenyon cells are determined by the signals from the projection neurons. The connections between the projection neurons and the Kenyon cells are fixed random connection [31] so that the outputs from the Kenyon-cells can be regarded as a random projection of sensory inputs. According to Johnson-Lindenstrauss Lemma [32][33], an appropriate random matrix $\boldsymbol{R}$ that satisfies following condition in a high probability exists.:

$$(1-\epsilon)\|\boldsymbol{x_1}-\boldsymbol{x_2}\|^2 \leq \|\boldsymbol{R}x_1-\boldsymbol{R}\boldsymbol{x_2}\|^2 \leq (1+\epsilon)\|\boldsymbol{x_1}-\boldsymbol{x_2}\|^2, \tag{29}$$

where $\boldsymbol{x_1}$and$\boldsymbol{x_2}$are arbitrary two different sensory inputs and $0<\epsilon<1$. (Note: $\dim(\boldsymbol{x}) > \dim(\boldsymbol{Rx})$ ). Thus, there are possibilities that Kenyon-cell outputs are represented as $Kenyon(\boldsymbol{x}) \simeq \boldsymbol{R}x$ and is also satisfying the same conditions as Eq 29. This means that the state changes in sensory input space $\boldsymbol{x}(t) \sim p(\boldsymbol{x}|\boldsymbol{x}(t-1), \boldsymbol{A}(t-1))$ can be converted to the state changes in the Kenyon-cell output space:

$$\boldsymbol{k}(t) \sim p(\boldsymbol{k}|\boldsymbol{k}(t-1), \boldsymbol{A}(t-1)). \tag{30}$$

This implies that the dynamics of $\boldsymbol{k}(t)$ can be approximately described by a Hidden Markov model.

#### Validity of analyzing Anti-Hebbian learning as Hebbian learning

MBON $\alpha'3$ compartment learns novel odor by anti-Hebbian learning. However, we constructed a simplified Hebbian learning model for the analysis. Here, we compare the reliabilities of anti-Hebbian and Hebbian learning in our model.

A novel detector can also be constructed using Hebbian learning. Therefore, the behavior of MBON $\alpha'3$ is also represented by active neurons, which are inactivated by a feature extractor formed by Hebbian learning. This implies that we can analyze the Hebbian learning part instead of the anti-Hebbian learning part. The $i$-th MBON neuron output is denoted as

$$u_i[\boldsymbol{x}] = f[\boldsymbol{W_i^T}\,\boldsymbol{x}], \tag{31}$$

where $\boldsymbol{W}_i$ is the weight vector of the $i$-th MBON neuron, and $f[a] = \max(a, 0)$ is the activation function. The input vector is denoted as $\boldsymbol{x_p} = [1, x_{p1}, \ldots, x_{pN}]^T$. Non-kernelized version of the model1,2 and 3 share the Hebbian learning rule with weight decay expressed as follows:

$$\Delta\boldsymbol{W_i} = \{-(1-\phi)W_i + \eta\boldsymbol{x_p}\}DAN[\boldsymbol{x_p}], \tag{32}$$

where $\phi < 1$ is the ratio of weight shrinking, $DAN[\boldsymbol{x}] \in \{0,1\}$ denotes activation of dopaminergic neuron. If the neuron receives several inputs $\boldsymbol{x_p}$ for $p = 1,2, \ldots,$ the weight vector converges in the following direction.

$$\boldsymbol{W_i} = \eta \left(\frac{1}{|\overline{S_D}|(1-\phi)}\right) \sum_{p \in \overline{S_D}} \boldsymbol{x_p}, \tag{33}$$

where $\overline{S_D}$ is the set of input indices that activates the dopaminergic neuron. This implies that weight vector $\boldsymbol{W}_i$ is the sum of the input vectors that activate the neuron. Note that if the number of inputs is large, the weight vector norm $\| \boldsymbol{W}_i \|$ also becomes large. Therefore, a weight decay term is necessary to avoid misestimation of the cos-similarity between the weight and input vectors. After convergence of the weight vector, the neuron output for $\boldsymbol{x}$ is expressed as follows:

$$u_i[x] = f\left[\left(\eta \sum_{p \in \bar{S}_i} \boldsymbol{x}_p\right)^T \boldsymbol{x}\right] = f\left[\eta \left(\frac{1}{|\overline{S_D}|(1-\phi)}\right) \left\|\sum_{p \in \bar{S}_D} \boldsymbol{x}_p\right\| \|\boldsymbol{x}\| \cos(\theta)\right], \tag{34}$$

where $\theta$ is angle between $\left(\frac{1}{|\overline{S_D}|(1-\phi)}\right) \sum_{p \in \overline{S_D}} \boldsymbol{x_p}$ and $\boldsymbol{x}$ . It is well known that each odor activates only 5 to 10 % of Kenyon cells [1], so that we can assume that $\boldsymbol{x}$ is a sparse vector and $\|\boldsymbol{x}\| \simeq constant$ and the learning samples are approximately satisfied that $\boldsymbol{x_p} \perp \boldsymbol{x_q}$ for $p \neq q$. Thus $u_i[x]$ is mainly dependent on $\cos(\theta)$, and $u_i$ is only activated when $\boldsymbol{x}$ is similar to one of $\boldsymbol{x}_p$ for $p \in \overline{S_D}$.

By contrast, for anti-Hebbian learning, the weight vector is updated as follows.

$$\Delta \boldsymbol{W}_i = \{(\boldsymbol{I} - \boldsymbol{W}_i)(1 - \phi) - \eta x_p\} DAN[\boldsymbol{x_p}] \tag{35}$$

converges in the following direction:

$$\boldsymbol{W}_i = \boldsymbol{I} - \eta \left(\frac{1}{|\overline{S_D}|(1-\phi)}\right) \sum_{p \in \overline{S_D}} \boldsymbol{x_p}, \tag{36}$$

where $\bar{S}_D$ is the set of indices of inputs that activate dopaminergic neuron, and $\boldsymbol{I}$ is the initial weight vector. After convergence of the weight vector, the neuron output for $\boldsymbol{x}$ is expressed as follows:

$$u_i[x] = f\left[\left(\boldsymbol{I^T x} - \eta \left(\frac{\mathbf{1}}{(1-\phi)|\overline{S_D}|}\right) \sum_{p \in \bar{S}_D} \boldsymbol{x}_p\right)^T \boldsymbol{x}\right] = f\left[\boldsymbol{I^T x} - \left(\frac{\mathbf{1}}{(1-\phi)|\overline{S_D}|}\right) \left\|\sum_{p \in \bar{S}_D} \boldsymbol{x}_p\right\| \|\boldsymbol{x}\| \cos(\theta)\right], \tag{37}$$

where $\theta$ is the angle between $\left(\frac{\mathbf{1}}{(1-\phi)|\overline{S_D}|}\right) \sum_{p \in \bar{S}_D} \boldsymbol{x}_p$ and $\boldsymbol{x}$. Let us assume that the initial weight vector $\boldsymbol{I} = [1,1,\dots,1]^T$. We can also assume that $I^T \boldsymbol{x}$ is constant for all inputs because we can assume that .each $\boldsymbol{x}$ is a sparse input vector and $\|\boldsymbol{x}\| \cong constant$. This means that the neuron output $u_i[\boldsymbol{x}]$ is activated only when $\boldsymbol{x}$ is not similar to any of $\boldsymbol{x}_p$ for $p \in \bar{S}_D$. This corresponds to the novelty detection. Notably, anti-Hebbian learning and Hebbian learning assign weights in the same direction but with opposite signs.

Therefore, we can analyze the anti-Hebbian learning part by analyzing the Hebbian learning part.

#### Feasibility of the three kernelized models as actual Hebbian learning models

Until now, the learning model of MBON $\alpha'3$ has been discussed as a kernelized model. However, we must consider whether the kernelized model is feasible as a true Hebbian learning model.

To examine this, we assumed that the output vectors of the Kenyon cells for different odors were orthogonal. These findings have been reported in several research papers [25] [26] [27]. Under this assumption, the learning samples are orthogonal.

The kernel method can be interpreted as the Hebbian learning model of a single neuron. In this section, this is referred to as the "target neuron."

We then demonstrate the feasibility of these three models. The activation condition of the target neuron is determined by the inner product of the input and weight vectors.

$$y(t) = f\left[\sum_{i=1}^{N} w_i(t) x_i(t)\right], \tag{38}$$

where $f[x]$ is assumed to be $max(0, x)$.

In particular, if each synaptic connection has its own $\sigma_i$, corresponding to the $\sigma_i$ value in lemma 2, then $\sigma_i$ is gradually reduced when the dopamine neuron is activated. Thus, the learning model of MBON $\alpha'3$ can be implemented as a true Hebbian learning model.

**Model 1:**

For Model 1, the synaptic weight is updated as follows. When a new sample is presented to the target neuron, if the inner product through the activation function (Eq. 39) is less than a certain threshold, and if $\sigma_i$ is less than a threshold value $\theta$, and the dopaminergic cell is activated,

$$w_i(t) = 0, \quad \sigma_i(t) = 1. \qquad (39)$$

This indicates that the synaptic weight is reset to 0 when dopaminergic neurons are activated. The target neuron learns the presented sample by using the following update rule:

$$w_i(t+1) = [w_i(t) + \eta x_i(t)]DAN[\boldsymbol{x}(t)], \qquad \sigma_i(t + 1) = 1 \qquad (40)$$

Here, $DAN[\boldsymbol{x}(t)]$ is the output of a dopaminergic neuron, and $DAN[\boldsymbol{x}(t)] = 1$ when the dopaminergic neuron is activated; otherwise, $DAN[\boldsymbol{x}(t)] = 0$ (see Figure S1). In this equation, we also assume that $w_i(t)$ is non-negative.

Eq. 39 corresponds to local vector elimination. Local vector elimination also corresponds to the removal of one kernel. Such local vector elimination can be realized using two possible mechanisms. One possibility is elimination by Bergmann glia (BG) nibbling parts of the synapses [28]. Specifically, it is thought to be realized by BG nibbling parts of synapses, which is triggered by local exposure (or externalization) of the "eat me" signal (phosphatidylserine [PS]) near less frequently used synapses. However, this occurs only in the brains of mice and not in those of Drosophila. A similar phenomenon is also observed in the Drosophila brain, but it occurs during a crucial period [29]. The most valuable possibility might be synapse reduction caused by the firing of the DAN in the absence of sensory inputs [30] [2].

Else, if $\sigma_i$ is larger than or equal to the threshold $\theta$, then Eqs. 39 and 40 are not executed. Instead, if the dopaminergic neuron is activated,

$$\sigma_i(t + 1) = \phi\sigma_i(t), \tag{41}$$

for all $i$.

**Model 2:**

For Model 2, the synaptic weight is updated as follows. When a new sample is presented to the neuron, if the dopaminergic neuron is activated, then Eqs. 39 and 40 are executed whenever $\sigma_i$ is one of the smallest values in the neuron. This differs from Model 1. Else, if $\sigma_i$ is not one of the smallest values and the neuron is activated, then Eq. 41 is executed.

**Model 3 (LRU):**

Model 3 is similar to Model 2, but the difference is that when a new sample is presented to the neuron, if the dopaminergic neuron is activated, Eqs. 39 and 40 are executed when $\sigma_i$ has the smallest value in the neuron.

If the dopaminergic neuron is activated, then Eq. 41 is executed. Meanwhile, the synapses with $x_i(t) > 0$ have their $\sigma_i$ reset to 1. The last process is added to reflect the least recently used (LRU) strategy.

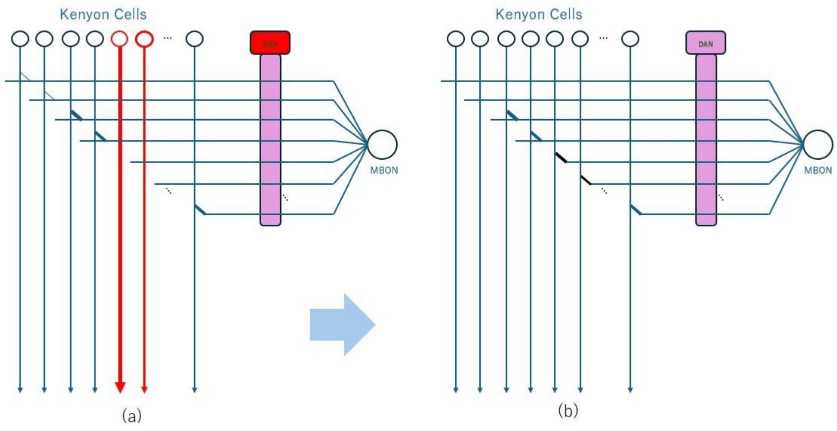


Fig. S1 An example of before:(a) and after:(b) snapshots of applying Eq 39 and Eq 40 in the case of Hebbian learning. The red colored Kenyon cells are activated cells and they represent the sparse feature vector of presented odor. Such weight changes are triggered by the activation of the DANs.
